# Agricultural Management Restructures the Functional Composition of Soil Fungal Communities Across Tropical Landscapes

**DOI:** 10.64898/2026.11.16.767036

**Authors:** Evelin Y. Reyes-Mendez, Cleber Ten Caten, Pedro Henrique Dias dos Santos, David Galo, Daijiang Li, Jordan A. Dowell

## Abstract

Soil fungal communities regulate key ecosystem processes, yet the extent to which agricultural management restructures fungal diversity across tropical landscapes remains poorly understood. We characterized fungal communities in 124 soil samples collected from natural vegetation, organic farms, and non-organic farms across four Honduran departments using internal transcribed spacer 2 (ITS2) and small-subunit ribosomal RNA (SSU) metabarcoding. Land use was associated with Bray-Curtis community composition after adjusting for geography, sequencing run, and read depth (ITS2 R2 = 0.06, SSU R2 = 0.05, both P < 0.001). Community dissimilarity increased with geographic distance within each land-use category (pooled Mantel r = 0.30 for ITS2 and 0.19 for SSU), and land-use categories shared 2.0-7.1 times as many fungal features as expected (all adjusted P < 0.001). For ITS2, total richness in non-organic soils was 18% lower than in natural soils and 25% lower than in organic soils. Across both markers, land-use restricted richness was 58-85% lower in non-organic soils than in natural and organic soils. In addition, symbiotrophs in non-organic soils were 18-20% of that in natural soils, while saprotroph and pathotroph responses varied by marker. Together, these findings demonstrate that agricultural management is associated with changes in fungal diversity, occupancy, and predicted trophic composition without eliminating the geographic structure or extensive taxonomic overlap of regional communities.

## Introduction

Soil fungi are fundamental drivers of terrestrial ecosystem function, mediating decomposition, nutrient cycling, soil aggregation, carbon storage, and plant–environment interactions. Through ecological strategies such as saprotrophy, symbiosis, and pathogenicity, fungi regulate energy and nutrient cycling while influencing plant productivity, community composition, and ecosystem resilience (Van Der Heijden et al. 1998; Powell and Rillig 2018). Shifts in fungal community composition can therefore alter ecosystem processes even when overall richness changes relatively little, because different ecological strategies perform distinct functions in soils (Wagg et al. 2014; Liu et al. 2023). Although global surveys have revealed strong geographic and environmental structure in soil fungal communities, substantial fungal diversity remains undescribed, particularly in tropical regions (Blackwell 2011; Hawksworth and Lücking 2017; Tedersoo et al. 2014, 2020; Mikryukov et al. 2023). As agricultural expansion intensifies across the tropics, understanding how fungal taxonomic and functional composition varies among land-use systems is therefore essential, rather than relying on richness alone as a measure of belowground biodiversity.

Community assembly offers a framework for interpreting this variation, as environmental filtering, dispersal, species interactions, and ecological drift influence which members of a regional species pool occur locally (Keddy 1992; Vellend 2010; Nemergut et al. 2013). Geographic structure can reflect dispersal opportunities and environmental differences among locations, whereas local vegetation and management may favor different taxa within a broadly shared assemblage. For example, among root-associated fungi, host plants, soil conditions, and dispersal constraints all contribute to community structure (Vályi et al. 2016; Horn et al. 2017), illustrating how regional and local filters can interact, and these same spatial and habitat associations are observed among broader soil fungal assemblages (Goldmann et al. 2016; Lumibao et al. 2020). However, while observational comparisons can identify these associations, identifying mechanisms of community assembly requires assessments beyond community composition.

Among anthropogenic influences on the environment, agricultural management changes several conditions relevant to fungal assembly. For instance, fertilization, pesticides, tillage, crop selection, and organic amendments can alter nutrient availability, carbon inputs, vegetation, and disturbance, leading to variation in soil biodiversity across agricultural systems (Bardgett and van der Putten 2014; Tsiafouli et al. 2015; Hartmann et al. 2015). Specifically, organic farming can support higher biodiversity on average than traditional farming practices, but the effects depend on land-use intensity and the organisms considered (Tuck et al. 2014). Further, host availability, in the form of crop diversity, can strongly influence individual groups of taxa, such as arbuscular mycorrhizal fungi, independently of a broad management label (Guzman et al. 2021). Thus, organic and conventional categories describe sets of practices whose ecological consequences may differ among landscapes.

Evidence of agriculture’s impacts on soil biodiversity in tropical systems remains less extensive than for temperate agriculture (Gumiere et al., 2016). Tropical land-use studies nevertheless document changes in fungal composition and guild representation following agricultural conversion (Brinkmann et al. 2019; Díaz-Vallejo et al. 2021). Such changes can occur within landscapes that differ in climate, vegetation, soils, and management history, limiting direct extrapolation between regions. In Central America, Honduras has natural vegetation, smallholder agriculture, and farms with contrasting management practices across environmentally heterogeneous departments, leading to large variations in soil properties and land-use classifications (Saravia-Maldonado et al. 2025). Honduras’ soil diversity serves as a mesocosm of heterogeneous environments, enabling comparisons of fungal communities across management categories while explicitly accounting for geographic structure.

We compared soil fungal communities from natural areas, organically managed farms, and non-organic farms across Intibucá, Olancho, Francisco Morazán, and El Paraíso using internal transcribed spacer 2 (ITS2) and small subunit ribosomal RNA (SSU) metabarcoding. We asked whether land use was associated with fungal richness, community composition, taxonomic read abundance, and predicted trophic modes after accounting for sequencing effort and spatial structure. We expected management categories to differ in composition and trophic representation, with lower symbiotroph representation in agricultural soils, while geographic turnover would persist within land uses.

## Materials and methods

### Study locations and soil sampling

We collected 124 soil samples between January and March 2024 across four departments in Honduras: Intibucá, Olancho, Francisco Morazán, and El Paraíso (Fig. 1; Table S1, Online Resource 1). We sampled natural areas and farms classified as organic or non-organic using interviews with owners or managers and field observations. Organic farms reported avoiding synthetic fertilizers, herbicides, and fungicides and using organic amendments, including composted manure and crop residues. Non-organic farms used synthetic fertilizers and pesticides. Natural areas comprised forest or other vegetation without obvious cultivation, fertilization, or pesticide application. Management categories were not independently certified or assigned experimentally.

**Fig. 1.**
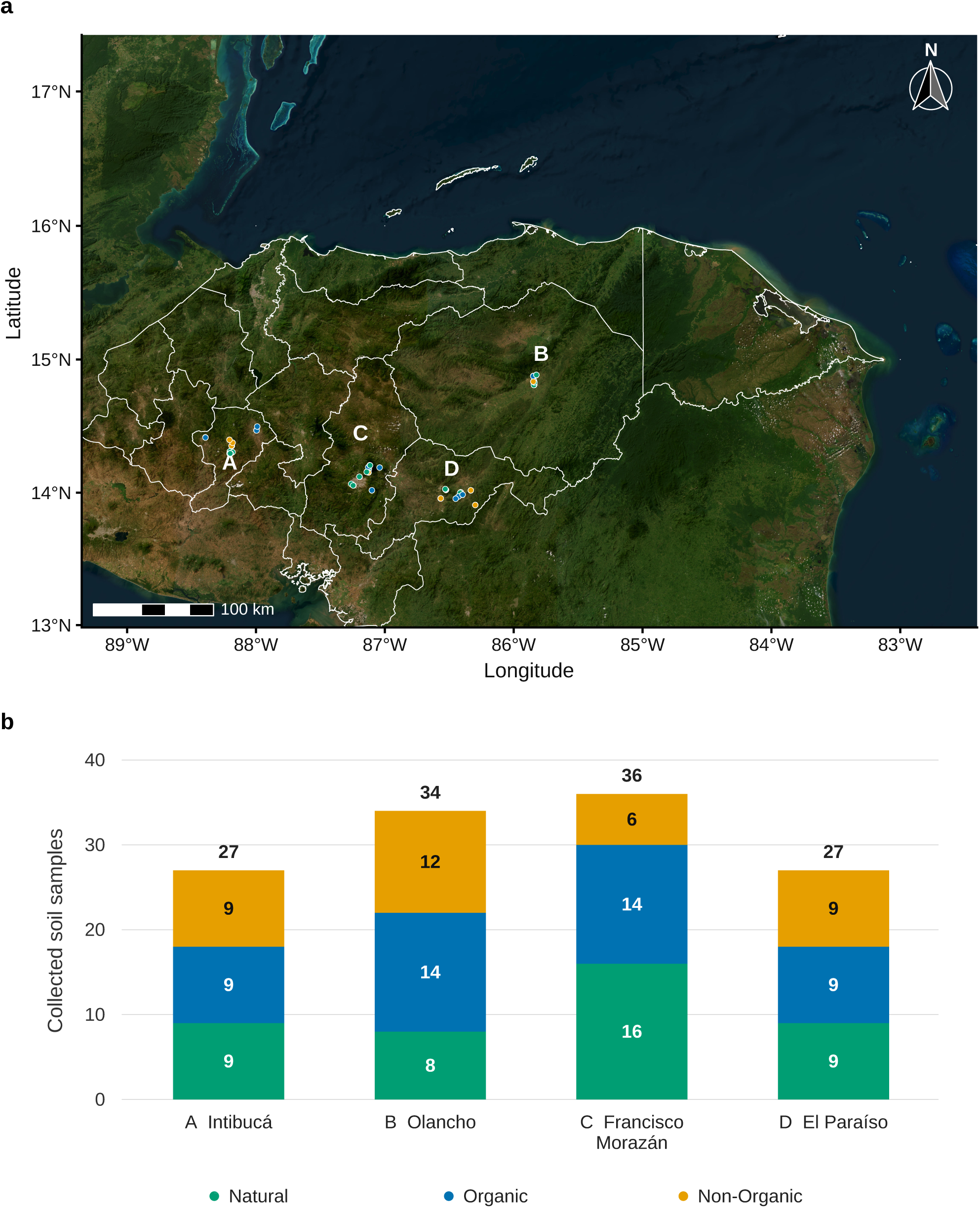
Sampling locations and land-use representation in Honduras. (a) Locations of the 124 collected soil samples. (b) Sample counts by department and land-use category. Segment labels give counts, and numbers above bars give department totals. Colors show natural (green), organic (blue), and non-organic (yellow) soils.

In 30 × 30 m plots, we removed surface litter and debris and collected three soil cores to a depth of 10 cm using a metal corer and rubber mallet. We cleaned the corer between samples by flaming, washing with 10% bleach, and rinsing with distilled water. Soil from the three cores was combined in a plastic bag and mixed by hand for at least 2 min. We transferred a composite subsample to a labeled microcentrifuge tube and stored it in a secondary bag with silica gel before laboratory processing. Geographic coordinates, collection dates, field labels, vegetation, and management classifications are recorded in Table S1.

### DNA extraction, sequencing, and bioinformatics

DNA was extracted at Escuela Panamericana El Zamorano, Honduras, using the DNeasy PowerSoil Pro Kit (QIAGEN, Hilden, Germany) according to the manufacturer’s instructions. We transported the extracted DNA in screw-cap tubes to Louisiana State University, USA, where we assessed its concentration and quality using a NanoDrop spectrophotometer (Thermo Fisher Scientific, Waltham, Massachusetts, USA). Samples were prepared in volumes of 50–500 µL at 400–600 ng/µL in 10× Tris–EDTA (TE) buffer.

We amplified the SSU rRNA gene using WANDA (5′-CAGCCGCGGTAATTCCAGCT-3′; Dumbrell et al. 2011) and AML2 (5′-GAACCCAAACACTTTGGTTTCC-3′; Lee et al. 2008), primers targeting arbuscular mycorrhizal fungi. ITS2 was amplified with ITS3 (5′-GCATCGATGAAGAACGCAGC-3′) and ITS4 (5′-TCCTCCGCTTATTGATATGC-3′; White et al. 1990). Paired-end amplicon sequencing was performed by The Scripps Research Institute, La Jolla, California, USA, on the AVITI platform (Element Biosciences, San Diego, California, USA). The SSU analyses below include all retained fungal features recovered by that assay, rather than treating the library as exclusively arbuscular mycorrhizal.

Reads were processed with LotuS 2.34, including quality filtering and chimera screening (Özkurt et al. 2022). ITS2 operational taxonomic units (OTUs) were clustered with VSEARCH at 97% identity (Rognes et al. 2016) and classified against UNITE v9 using BLAST-based lowest-common-ancestor assignment (Abarenkov et al. 2023, 2024). SSU amplicon sequence variants (ASVs) were inferred with DADA2 within LotuS (Callahan et al. 2016). SSU classification combined a custom virtual-taxon reference with a SILVA 138.1 reference file designated as non-arbuscular-mycorrhizal (Quast et al. 2013). The logged commands, reference basenames, quality-filter configurations, and marker-specific processing distinctions are provided in Note S1 (Online Resource 2).

### Marker-level community analyses

We analyzed ITS2 OTU and SSU ASV datasets separately (due to differences in primer affinity) using a common framework in R 4.6.0, with phyloseq and vegan for data handling and community analyses (McMurdie and Holmes 2013; Oksanen et al. 2026). We retained features assigned to *Fungi* or recognized fungal phyla. We selected minimum fungal read depths from individual-sample rarefaction curves, estimated sample coverage, and retention across land uses. We evaluated cutoffs from 250 to 20,000 reads and selected the lowest cutoff with median coverage of at least 95% and 10th-percentile coverage of at least 90% across all land-use categories. This yielded 3,000 reads for ITS2 and 2,000 for SSU (Fig. S1, Online Resource 3; Table S1). Coverage describes estimated read-probability mass rather than the fraction of taxa recovered. Because richness curves continued to accumulate taxa, primary models used unrarefied data with explicit depth adjustment (Tedersoo et al. 2022).

We compared read depths among land uses with Kruskal-Wallis tests and examined richness-depth relationships using linear models with log_10_ read depth, land use, and their interaction (Figs. S2 and S3). Primary community analyses used Bray-Curtis dissimilarities of relative fungal read abundances and principal coordinates ordination analysis (Fig. S4). Haversine distances among sampling coordinates were summarized by two geographic principal coordinate axes. We tested marginal land-use effects with permutational multivariate analysis of variance (PERMANOVA; Anderson 2001), adjusting for these axes, sequencing run, and log_10_ fungal read depth. We used 9,999 unrestricted permutations and repeated tests with permutations restricted within recorded sampling locations. Multivariate dispersion used group-size bias adjustment, permutation tests, and Tukey-adjusted pairwise comparisons (Anderson 2006; Fig. S5; Table S2). In addition, geographic distance decay was evaluated with Spearman Mantel tests and 9,999 permutations, pooled and within land uses (Fig. S6). Community sensitivity analyses used Jaccard presence-absence, Hellinger-transformed Euclidean, and Aitchison distances. For Aitchison analyses, we retained features present in at least 5% of samples, removed empty samples or those with more than 99% zeros, applied count-zero multiplicative replacement (Palarea-Albaladejo and Martín-Fernández 2015), and calculated Euclidean distances of centered log-ratio values. We also repeated Bray-Curtis and richness analyses at a stricter 5,000-read cutoff (Fig. S7; Table S3). We report unrestricted and within-location tests separately and interpret composition results alongside dispersion.

### Richness and fungal community structure

For community overlap, features were retained within a land-use category if present in at least 10% of its samples. We tested intersections against the set-size-conditioned null using SuperExactTest (Wang et al. 2015) and false-discovery-rate (FDR) correction (Table S4). Separately, we partitioned each sample’s complete observed richness into features detected in more than one land-use category and features detected only in one category. The latter describes observed sampling occupancy, not demonstrated ecological exclusivity.

We modeled total, shared, and land-use-restricted richness using spatial negative binomial models that included land use, land-use-specific log_10_ read-depth slopes, sequencing run, and a Matern covariance structure across sample coordinates. Read depth was centered at the primary-cohort median on the log_10_ scale. Pairwise contrasts were evaluated at this reference depth, which was held constant in the 5,000-read sensitivity analysis (Fig. S8; Table S5). Secondary richness comparisons used deterministic expected rarefied richness at the dataset read-depth tenth percentile. Pooled land-use assemblages were additionally standardized to the smaller of 90% coverage and the minimum observed assemblage coverage using iNEXT (Chao et al. 2014; Hsieh et al. 2016). We distinguished these pooled estimates from sample-level richness, testing overall richness effects using likelihood-ratio comparisons against reduced models lacking the land-use intercept and depth-interaction terms, yielding joint four-degree-of-freedom tests.

We aggregated fungal reads by phylum, class, and predicted FUNGuild trophic mode. Only Probable or Highly Probable annotations were retained (Nguyen et al. 2016). Analysis of trophic mode groups required at least five positive samples in every land-use category and at least 10% overall prevalence for count-model inference. Spatial negative-binomial abundance models were fitted with spaMM (Rousset and Ferdy 2014) and included land use, sequencing run, a log fungal-read-depth offset, and Matern covariance (Tables S6 and S7). We checked multiple optimization starts and full/null likelihood ordering and derived all contrasts from a single full fit. Benjamini– Hochberg correction (Benjamini and Hochberg 1995) was applied separately by marker, cutoff, and response rank across overall tests or all pairwise contrasts. Overall abundance tests compared full and reduced models differing in the two land-use coefficients. We use q to denote Benjamini– Hochberg-adjusted P values.

## Results

### Community composition and geographic turnover

Of the 124 collected samples, the selected cutoffs retained 111 ITS2 samples (37 natural, 43 organic, and 31 non-organic) and 92 SSU samples (34 natural, 34 organic, and 24 non-organic) (Fig. 1, Table S1). Retained groups had median estimated sample coverage above 95%, although richness curves did not fully plateau (Fig. S1). Read-depth differences among land uses were not significant for either marker (Fig. S2; Table S1), and richness remained positively related to read depth (Fig. S3; Table S5). Land use was associated with Bray-Curtis community composition after adjustment for geography, sequencing run, and read depth (ITS2: R² = 0.06; SSU: R² = 0.05; both P < 0.001). The ITS2 effect remained significant within sampling locations (P < 0.001), whereas the SSU within-location test did not (P = 0.09; Table S2). Communities shared more features than expected under the set-intersection null (enrichment range 2.0-7.1-fold; all q < 0.001; Fig. 2; Table S4), despite differences in composition. In addition, Bray-Curtis dispersion differed among land uses for ITS2 (P < 0.001), but evidence was weaker for SSU (P = 0.15; Fig. S5). These dispersion differences indicate that PERMANOVA effects are shifts in centroids. Further, geographic associations were positive in both pooled datasets (Mantel r = 0.30 and 0.19, respectively; both P < 0.001), and within all land-use categories (Fig. 3 and Fig. S6; Table S2).

**Fig. 2.**
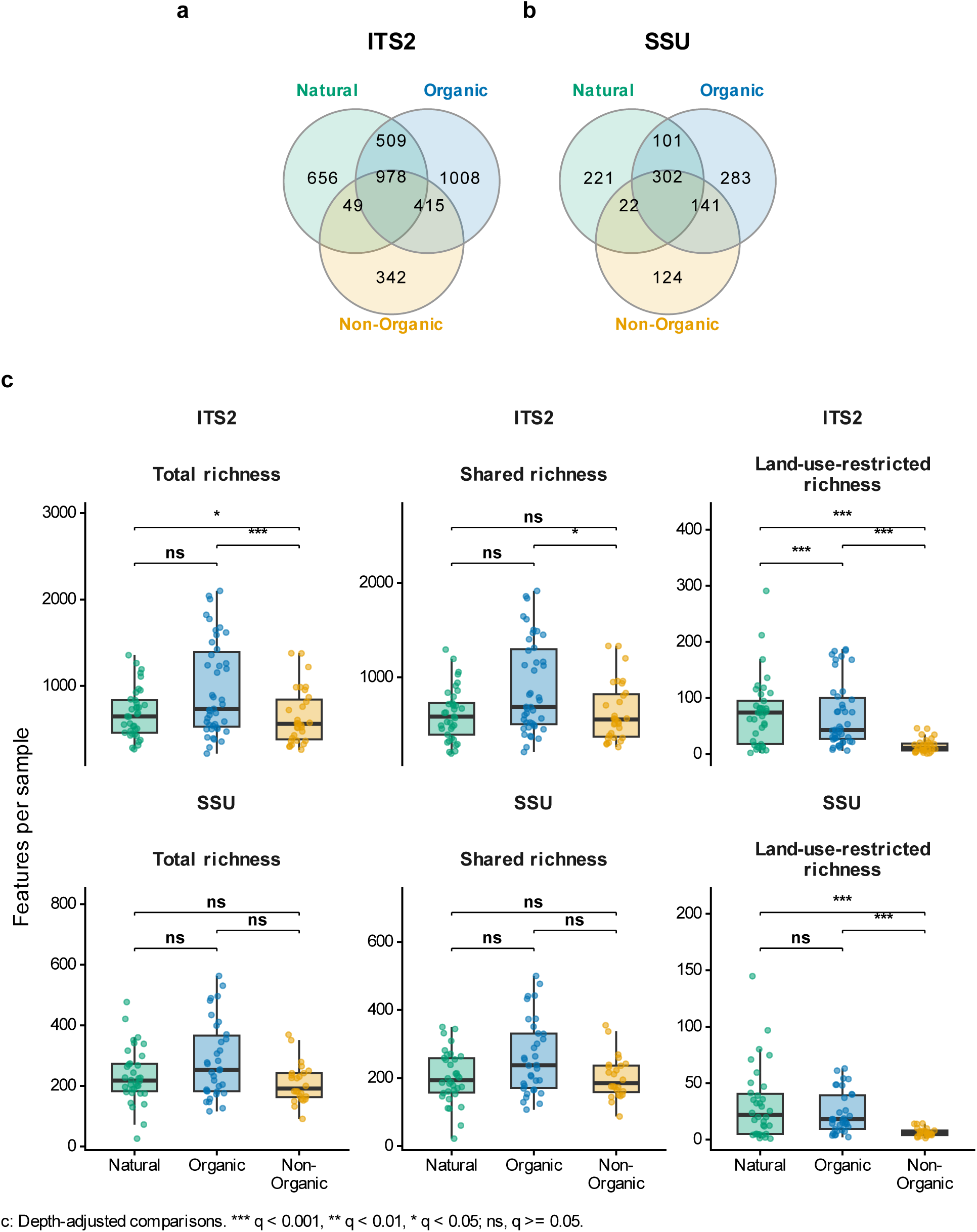
(a, b) Venn diagrams for ITS2 and SSU, respectively, and (c) sample richness across land uses. Venn diagrams show features detected in at least 10% of samples within each land-use category for ITS2 and SSU at their selected read cutoffs. The richness panels show total, shared, and observed land-use-restricted richness per retained sample. Shared features occur in more than one land-use category; restricted features were detected in only one. Points are individual samples, and boxplots show medians, interquartile ranges, and whiskers extending to 1.5 times the interquartile range. In panel c, brackets show pairwise contrasts from spatial negative-binomial richness models at the primary marker-specific reference read depth, with sequencing-run and spatial adjustment. Stars use Benjamini-Hochberg correction across all nine richness contrasts within each marker (* q < 0.05, ** q < 0.01, *** q < 0.001; ns, q ≥ 0.05). Boxes and points show unadjusted observations; the brackets summarize adjusted comparisons at the reference depth. Exact ratios, confidence intervals, and adjusted P values are reported in Table S5. Colors show natural (green), organic (blue), and non-organic (yellow) soils.

**Fig. 3.**
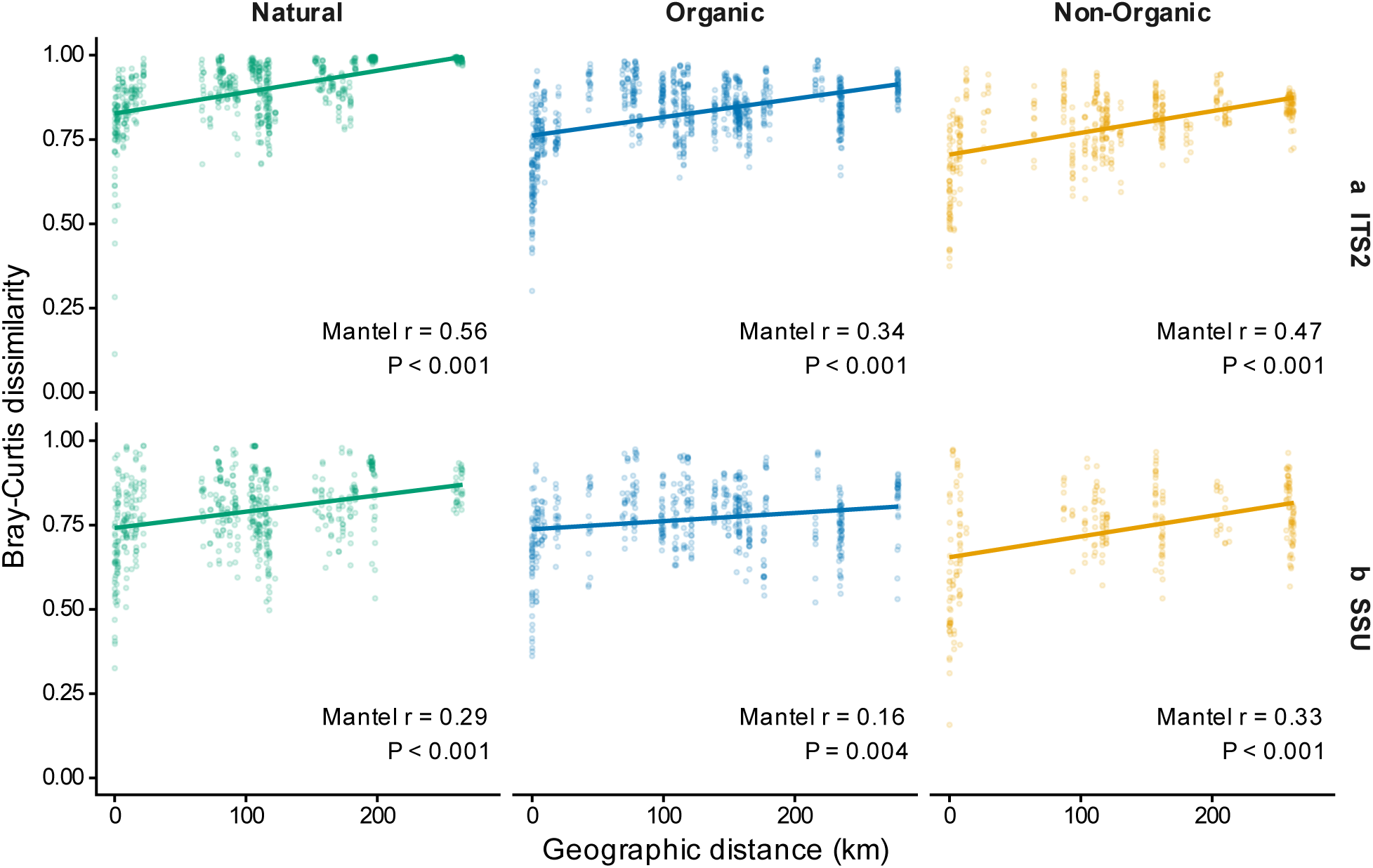
Geographic distance-decay within land-use categories. Pairwise Bray-Curtis dissimilarity is plotted against Haversine geographic distance, separately for each marker and land use. Lines are descriptive linear trends. Labels report Spearman Mantel correlations and permutation P values from 9,999 permutations. Pairwise points are dependent comparisons, not independent replicates. Table S2 reports the tests. Marker rows are (a) ITS2 and (b) SSU. Colors show natural (green), organic (blue), and non-organic (yellow) soils.

ITS2 mean observed unadjusted richness was 965.4 ± 545.2 SD OTUs in organic soils, compared with 682.6 ± 284.2 SD OTUs in natural soils and 642.5 ± 316.2 SD OTUs in non-organic soils. SSU mean observed unadjusted richness was 285.6 ± 126.8 SD ASVs in organic soils, compared with 233.5 ± 92.6 in natural soils and 208.4 ± 65.6 in non-organic soils (Fig. 2; Table S5). At the common ITS2 reference sequencing depth, non-organic soils had lower adjusted total richness than natural soils (ratio = 0.82, q = 0.02) and organic soils (ratio = 0.75, q < 0.001); however, the organic-versus-natural contrast was not significant (ratio = 1.09, q = 0.25). For SSU, joint land-use and depth-slope terms were significant for total and shared richness, but their individual category contrasts at the reference sequencing depth did not meet the 0.05 FDR threshold (Table S5). While observed richness was shared across land uses (Fig. 2), ITS2 non-organic soils had lower adjusted land-use-restricted richness than natural and organic soils (ratios = 0.32 and 0.15, respectively; both q < 0.001). SSU non-organic soils had lower adjusted land-use-restricted richness than natural and organic soils (ratios = 0.42 and 0.28, respectively; both q < 0.001). ITS2 organic soils had greater adjusted restricted richness than natural soils (ratio = 2.07, q < 0.001). Adjusted shared-richness contrasts were weaker overall, although ITS2 shared richness was lower in non-organic than in organic soils (ratio = 0.77, q < 0.05).

### Taxonomic and trophic structure variation

ITS2 *Basidiomycota* had lower read abundance in non-organic than in natural and organic soils (ratios = 0.32 and 0.33, respectively; both q < 0.05). SSU *Basidiomycota* had lower read abundance in non-organic soils than in natural and organic soils (ratios = 0.37 and 0.32, respectively; both q < 0.05). ITS2 *Mucoromycota* was lower in non-organic than in natural soils (ratio = 0.25, q < 0.05), but its non-organic-versus-organic contrast was not significant. However, for SSU, *Mucoromycota* showed the same pattern as *Basidiomycota* (ratios = 0.51 and 0.48; both q < 0.05) (Fig. 4). Class-level overall tests identified land-use associations in 6 of 36 classes for ITS2 and 2 of 20 modeled classes for SSU. Specifically, SSU *Archaeorhizomycetes* was lower in non-organic than in natural soils (ratio = 0.004, q < 0.001). SSU *Sordariomycetes* was higher in organic than in natural soils (ratio = 2.09, q < 0.05) and lower in non-organic than in organic soils (ratio = 0.51, q < 0.05). ITS2 *Sordariomycetes* did not show significant corrected pairwise differences. Full phylum and class results, including sparse-group screening, are reported in Tables S6 and S7 and Fig. S10.

**Fig. 4.**
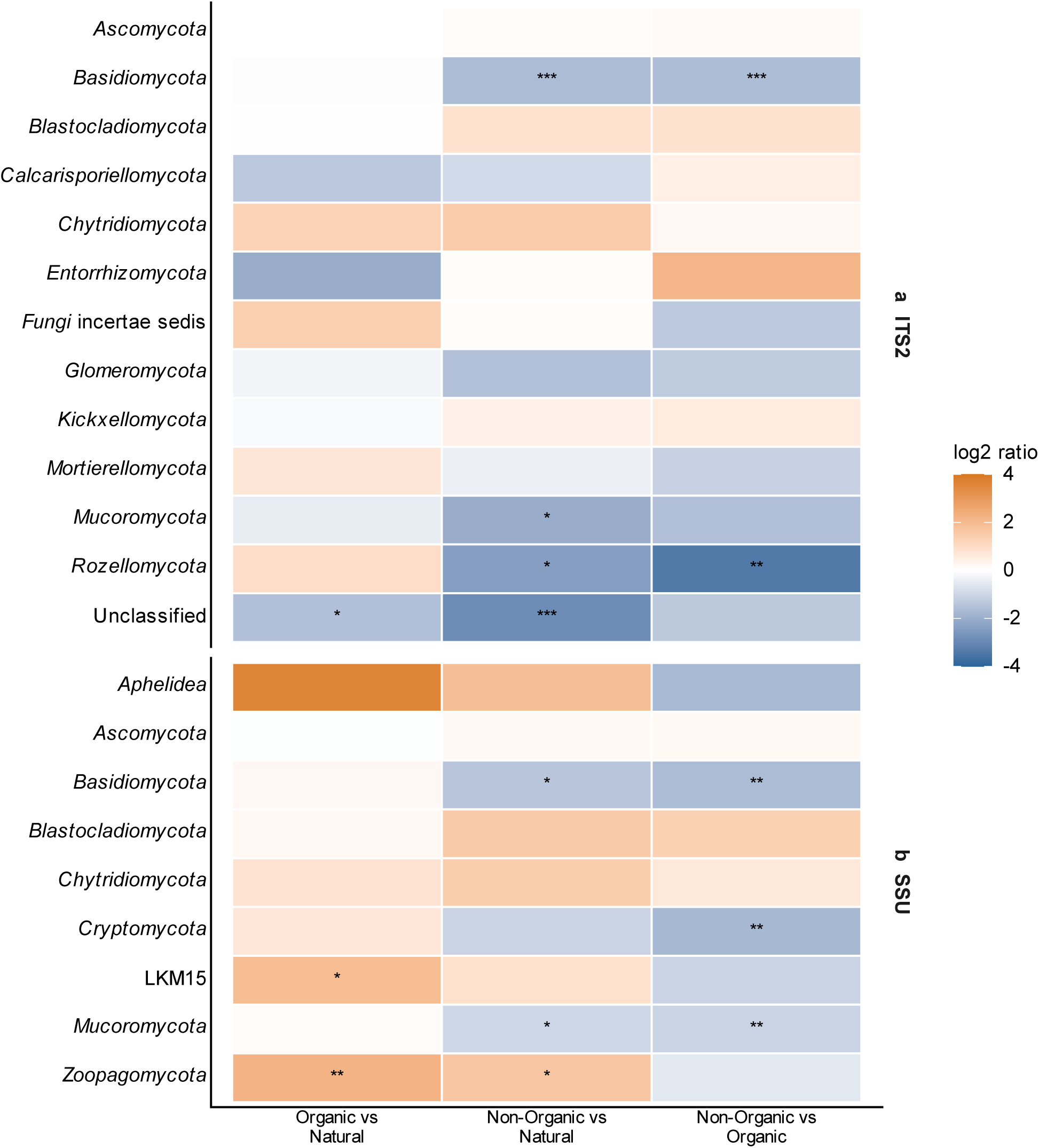
Land-use contrasts in phylum-level read abundance. Colors show model-estimated log₂ ratios for organic versus natural, non-organic versus natural, and non-organic versus organic soils. Ratios derive from a single spatial negative-binomial full model per group, with a fungal-read-depth offset, sequencing-run covariate, and geographic covariance. Color scales are capped at ±4. Stars indicate Benjamini-Hochberg-adjusted significance across all phylum contrasts within each marker (* q < 0.05, ** q < 0.01, *** q < 0.001). Marker rows are (a) ITS2 and (b) SSU

Among trophic-mode contrasts, pure symbiotroph read abundance was lower in non-organic than in natural soils for both markers (ITS2 ratio = 0.18, q < 0.001; SSU ratio = 0.20, q < 0.01). ITS2 pure symbiotrophs were also lower in organic than natural soils (ratio = 0.28, q < 0.01). ITS2 pure saprotrophs were higher in organic than natural soils (ratio = 1.49, q < 0.05), while SSU pure pathotrophs were higher in non-organic than natural soils (ratio = 7.55, q < 0.001). SSU pure saprotroph contrasts did not meet the 0.05 FDR threshold. Thus, guild responses differed between markers, with the clearest shared pattern involving lower symbiotroph representation in non-organic soils (Fig. 5; Tables S6 and S7).

**Fig. 5.**
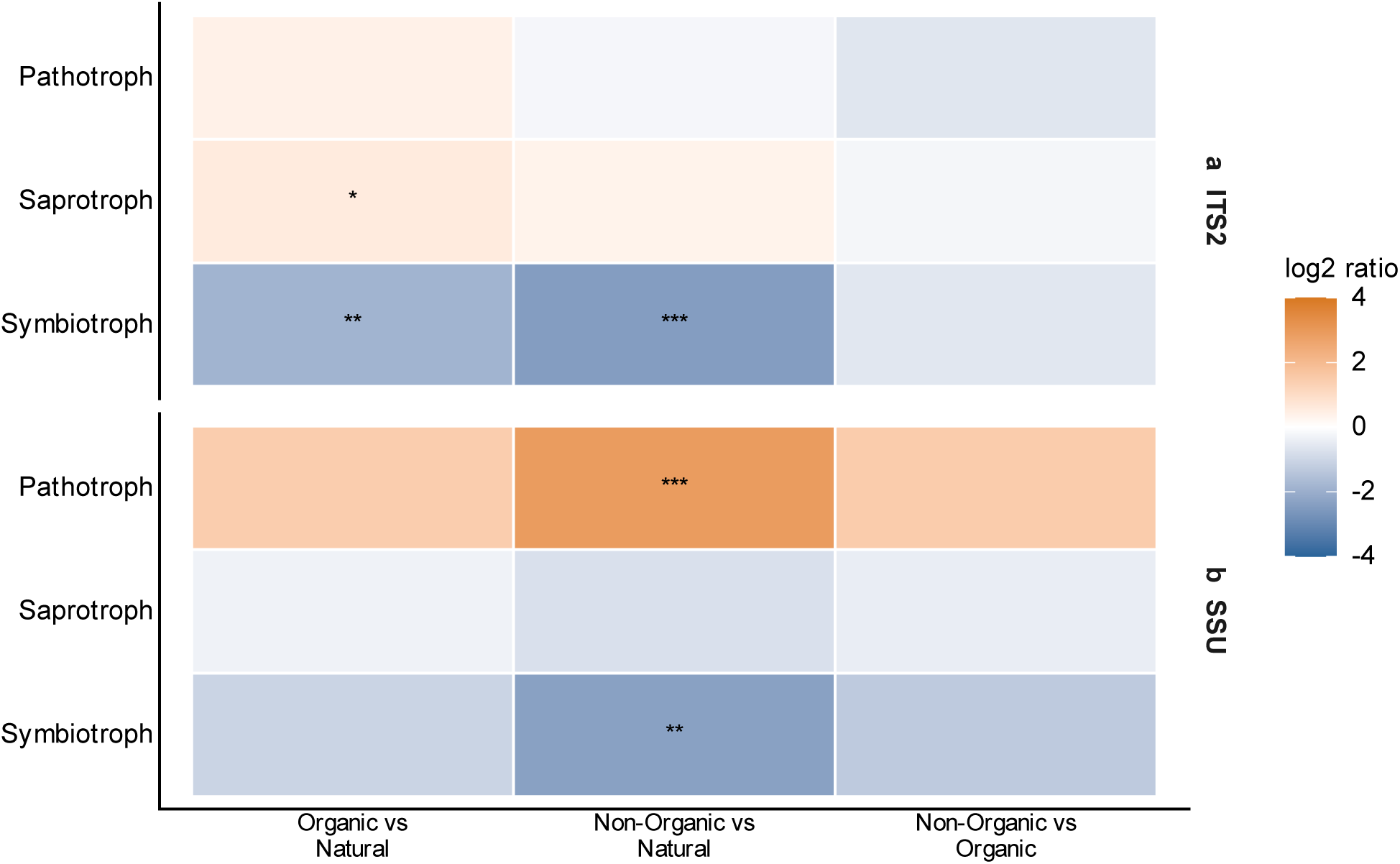
Land-use contrasts in the three pure trophic-mode categories: Pathotroph, Saprotroph, and Symbiotroph. Colors show model-estimated log₂ ratios for organic versus natural, non-organic versus natural, and non-organic versus organic soils. Ratios derive from a single spatial negative-binomial full model per group, with a fungal-read-depth offset, sequencing-run covariate, and geographic covariance. Color scales are capped at ±4. Stars indicate Benjamini-Hochberg-adjusted significance across all trophic-mode contrasts within each marker (* q < 0.05, ** q < 0.01, *** q < 0.001).

### Sensitivity analyses

The 5,000-read sensitivity cutoff retained 106 ITS2 and 59 SSU samples. ITS2 composition effects remained significant in unrestricted and within-location tests (P < 0.001 for the latter). SSU composition remained significant without site restriction, but the stricter within-location cutoff yielded P = 0.06. ITS2 effects also persisted across Jaccard, Hellinger, and Aitchison distances. SSU within-location results were significant for Jaccard (P < 0.01) and Aitchison (P < 0.001), but not Hellinger (P = 0.20; Fig. S7; Table S3). Richness effect sizes and uncertainty changed with filtering (Fig. S8; Table S5), so results were not treated as uniformly robust across every framework. In addition, dispersion remained metric-dependent (Table S3).

## Discussion

Land-use change in tropical landscapes is often framed as a progressive loss of biodiversity. However, for belowground biodiversity, the Honduran communities suggest a more nuanced transition. Specifically, natural areas and farms retained much of the same detected fungal assemblage but differed in the distribution of its members and in the representation of predicted symbiotic lifestyles. This distinction places our findings within a growing body of work showing that agricultural conversion can strongly alter fungal composition and guild structure even when total richness changes weakly or inconsistently. For example, tropical studies have reported substantial compositional shifts following conversion to plantations and croplands, yet the direction and magnitude of alpha-diversity responses vary among systems (Brinkmann et al. 2019; Song et al. 2019; Berkelmann et al. 2020; Díaz-Vallejo et al. 2021). Our Honduran results similarly indicate that richness alone can understate potentially ecologically meaningful differences among land uses.

### Land use reorganizes a shared fungal assemblage

The persistence of distance–decay across all land-use categories indicates that agricultural management did not erase the regional organization of fungal communities. Geographic structure is pervasive in fungi and can arise from dispersal limitation together with spatial variation in climate, vegetation, soils, and host availability (Bahram et al. 2013; Tedersoo et al. 2014, 2020; Mikryukov et al. 2023). In Honduras, land-use differences within a broadly shared fungal assemblage further suggest that regional processes strongly shape the pool of fungi encountered across the landscape. Similar distinctions between widespread “core” taxa and taxa associated with particular sites or land-use contexts have been reported in other agricultural systems (Gschwend et al. 2022). In the context of community assembly theory, regional dispersal and environmental history influence which fungi are available to occur at a site, whereas local vegetation, soils, management, species interactions, and ecological drift influence their local occurrence and relative representation (Keddy 1992; Vellend 2010; Nemergut et al. 2013). Although our observational data cannot distinguish among these mechanisms, they provide plausible explanations for how land-use-associated community differences can arise without forming entirely distinct fungal species pools that can be assessed more thoroughly.

Large-scale studies suggest that common and uncommon fungi can respond differently to agricultural environments, providing context for observations of lower land-use-restricted richness in non-organic soils. Abundant fungi often occupy broader environmental ranges than rare fungi in agricultural fields (Jiao and Lu 2020), while European arable soils contain fewer rare fungi and more widespread taxa than grasslands (Banerjee et al. 2024). These studies suggest that agricultural environments disproportionately constrain fungi with narrower distributions or environmental tolerances. However, we defined our land-use-restricted features by observed occupancy among categories rather than by abundance, prevalence, or niche breadth. They therefore cannot be identified directly as rare or specialized. The supported conclusion is narrower but still biologically meaningful, in that non-organic soils contained a smaller category-restricted component of the detected assemblage. Whether this pattern reflects environmental filtering, lower habitat heterogeneity, host differences, dispersal, or incomplete detection remains unclear.

The organic communities further demonstrate why land-use responses cannot be reduced to a simple continuum from natural to increasingly intensive agriculture. Organic soils supported high observed richness and a comparatively large ITS2 land-use-restricted component similar to natural areas, yet their communities did not reproduce the trophic profile of natural areas. This interpretation is consistent with long-term farming-system studies showing that organic and conventional management can generate compositionally distinct microbial communities rather than one system simply retaining more of the original community (Hartmann et al. 2015). Similarly, van Rijssel et al. (2022) found that fungal communities became increasingly distinct during conversion to organic management, but apparent increases in diversity disappeared when nearby conventional reference fields were included, emphasizing that management-attributed differences can depend strongly on paired environmental controls.

Broadly, the biological effects of organic management depend on the practices and environmental conditions encompassed by that label. Organic farming supports greater biodiversity on average, but the effect varies with land-use intensity and organismal group (Tuck et al. 2014). Specifically, crop diversity can enrich arbuscular mycorrhizal fungal communities even within an otherwise intensive landscape (Guzman et al. 2021), while amendments, vegetation, soil chemistry, and management duration can each influence fungal community composition independently of certification category (Alguacil et al. 2014; Hartmann et al. 2015; Manoharan et al. 2017). Thus, the organic communities in Honduras are best understood as distinct assemblages that may arise from combinations of crops, amendments, vegetation, and site histories, rather than as restored natural communities.

### Plant–fungal associations shift, but responses are selective

The most consequential biological pattern was the lower representation of fungi classified exclusively as symbiotrophs in non-organic soils. Its importance lies less in demonstrating a general decline of fungi than in suggesting that land use can change which ecological relationships are represented within communities that still share many members. Specifically, organic soils also had lower ITS2 pure-symbiotroph representation than natural soils, reinforcing the conclusion that high fungal richness does not necessarily preserve the trophic profile associated with natural vegetation. Other studies similarly show that plant-associated fungal groups can respond more strongly than aggregate richness. Tropical forest conversion to rubber plantation reduced ectomycorrhizal fungi despite little change in alpha diversity (Song et al. 2019). In a 24-year boreal field experiment, forests had the highest representation of symbiotrophs, whereas organic and conventional crop rotations had higher proportions of pathotrophs and plant pathogens; nevertheless, organic and conventional systems did not consistently differ in the potentially beneficial fungal groups examined (Häkkinen et al. 2025). Most recently, paired sampling across elevational and climatic gradients in Ecuador found substantially lower arbuscular mycorrhizal fungal richness in cultivated than in adjacent uncultivated soils, with the magnitude and direction of diversity relationships varying with temperature and precipitation (Stewart et al. 2026). Overall, agricultural effects on plant-associated fungi can be substantial but strongly depend on host vegetation, management, climate, soil conditions, and the fungal group examined.

The broader mycorrhizal literature cautions against treating symbiotic fungi as a single ecological unit, as variation in mycorrhizal diversity and community composition can shape nutrient acquisition, plant productivity, soil aggregation, ecosystem stability, and broader ecosystem multifunctionality even when richness alone changes little (van der Heijden et al. 1998; Wagg et al. 2014; Bender et al. 2016; Powell and Rillig 2018). For example, across ten Australian biomes, agricultural land use favored putative Mucoromycotinian arbuscular mycorrhizal fungi but not Glomeromycotinian arbuscular mycorrhizal fungi, and the responses of both groups depended on climate, soil pH, and plant richness (Albornoz et al. 2022). Management practices can similarly alter AMF composition without producing uniform changes in abundance or richness (Alguacil et al. 2014; Manoharan et al. 2017), and crop diversity can increase AMF diversity within agricultural landscapes (Guzman et al. 2021). This context helps explain why the Honduran taxonomic responses were selective rather than coordinated across every lineage that may contain plant-associated fungi. Consistent with previous results, the lower representation of Basidiomycota and Mucoromycota in non-organic soils and the strong SSU response of Archaeorhizomycetes identify particular land-use associations, but they do not demonstrate a general collapse of root-associated fungal diversity.

The marker-dependent saprotroph and pathotroph responses further argue against interpreting the communities as undergoing a uniform transition from mutualistic to disturbance-adapted lifestyles. ITS2 supported greater representation of pure saprotrophs in organic soils, whereas SSU supported greater representation of pure pathotrophs in non-organic soils. Neither response was independently reproduced by the other marker. This disagreement is not unexpected because ITS2 and SSU differ in primer specificity, taxonomic coverage, reference databases, and resolution (White et al. 1990, Lee et al. 2008, Dumbrell et al. 2011, Quast et al. 2013). The SSU assay targeted AMF but included all retained fungal ASVs, whereas ITS2 provided a broader community survey. The convergence on lower pure-symbiotroph representation is therefore notable, but the remaining guild responses should be interpreted as marker-specific evidence rather than components of a single coordinated ecological transition.

### Implications for tropical agroecosystems

Our Honduran results contribute to an increasingly diverse picture of fungal responses to tropical agriculture, where some tropical conversion studies have found major shifts in community composition and predicted guilds (Brinkmann et al. 2019; Berkelmann et al. 2020), while broader synthesis indicates that responses vary substantially among land uses, microbial groups, environmental contexts, and analytical approaches (Díaz-Vallejo et al. 2021). Tropical agricultural landscapes encompass steep climatic gradients, diverse soils, different crop hosts, contrasting land-use histories, and management systems ranging from smallholder polycultures to industrial monocultures. These factors can alter both the baseline regional assemblage and how fungi respond to cultivation. In this broader context, our Honduran study’s main contribution is to separate several dimensions of fungal change that are often combined. Agricultural communities retained substantial membership overlap with natural areas, yet non-organic soils had a smaller land-use-restricted component and markedly lower representation of pure symbiotrophs. Organic soils supported higher observed richness and greater ITS2-restricted richness but did not reproduce the natural-area trophic profile. These patterns show that neither total richness nor a broad management label is sufficient to characterize a fungal community’s ecological state. Composition, observed occupancy, trophic representation, and regional turnover reveal different aspects of the response.

This conclusion has practical implications, but the observational design does not justify prescribing particular farming practices. The organic and non-organic categories encompassed differences in crop identity, vegetation, amendments, synthetic inputs, disturbance, and management history. Studies using paired fields or long-term experiments show that controlling this background variation can substantially change the apparent effect of management (Hartmann et al. 2015; van Rijssel et al. 2022; Häkkinen et al. 2025). Future work in Honduras should therefore combine matched natural–agricultural comparisons and repeated sampling with direct measurements of soil chemistry, crop and non-crop plant diversity, tillage, fertilizers, pesticides, amendments, and management duration. Pairing these variables with root colonization, fungal biomass, nutrient acquisition, soil aggregation, pathogen incidence, and plant performance would establish whether the compositional patterns detected here translate into altered ecological processes. Further, incorporating metagenomic or metatranscriptomic sequencing would allow specific dissection of taxonomic vs functional diversity.

Overall, land use in Honduras was associated not with the explicit disappearance of a regional fungal biota, but with changes in how that biota was distributed and represented ecologically across management contexts. The strongest evidence concerned a contraction of the land-use-restricted component in non-organic soils and lower representation of predicted symbiotrophs, while geography continued to structure communities across the landscape. Viewed alongside the wider literature, these findings support a shift away from asking only whether agriculture reduces fungal richness and toward asking which community members and ecological associations are retained, redistributed, or diminished under specific environmental and management conditions.

## Supporting information

Electronic Supplemental Resource 1

Electronic Supplemental Resource 2

Electronic Supplemental Resource 3

## Acknowledgements

We thank Valeria Reyes, Wesly Ramirez, Carlos Valladares, Javier Bu, and Alicia Cruz for assistance with soil sampling and field logistics. Carolina Avellaneda of Escuela Panamericana El Zamorano provided laboratory facilities and equipment for DNA extraction. We thank the farmers, landowners, collaborators, and community members who provided access to sampling locations, and Michael Hellberg and Laura Lagomarsino for comments on the manuscript. OpenAI Codex assisted with generating figures.

## Statements and Declarations Funding

The Society for the Protection of Underground Networks supported this work through its Underground Explorers Program, awarded to Evelin Y. Reyes-Mendez, and the Department of Biological Sciences at Louisiana State University provided institutional start-up support to Jordan A. Dowell and Daijiang Li.

## Competing interests

The authors declare no competing interests.

## Author contributions

E.Y.R., C.T.C., and D.L. designed the study; E.Y.R. and C.T.C. collected the data; D. G., E.Y.R., and P.S. extracted the DNA samples and prepared them for sequencing. J.A.D. analyzed the data with input from E.Y.R., P.S and C.T.C.; and E.Y.R. and J.A.D. wrote the manuscript with input from all authors. All authors approved the final version of the manuscript.

## Data availability

Sequence data are associated with NCBI BioProject PRJNA1484736. Sampling metadata and numerical results are provided in Online Resource 1, bioinformatic methods in Online Resource 2, and supplementary figures in Online Resource 3.

## ESM 1: Supplementary tables

**Table S1.** Sampling metadata, retention, and read-depth selection

**Table S2.** Community composition, geographic turnover, and dispersion

**Table S3.** Community sensitivity analyses

**Table S4.** Community overlap and FUNGuild annotations

**Table S5.** Richness summaries, models, and sensitivity analyses

**Table S6.** Taxonomic and trophic-mode models and eligibility

**Table S7.** Taxonomic and trophic-mode pairwise comparisons

**Table S8.** Model diagnostics and coefficients

## ESM 2: Supplementary note

**Supplemental Note S1.** Bioinformatic processing of ITS2 and SSU libraries

## ESM 3: Supplementary figures

**Figure S1.** Rarefaction curves and read-cutoff selection.

**Figure S2.** Fungal sequencing depths of retained samples across land uses.

**Figure S3.** Observed richness in relation to fungal sequencing depth.

**Figure S4.** Principal coordinates ordination of retained fungal communities.

**Figure S5.** Within-land-use community dispersion.

**Figure S6.** Geographic distance-decay across all retained samples.

**Figure S7.** Sensitivity of community-composition results to distance metric and read cutoff.

**Figure S8.** Sensitivity of adjusted richness contrasts to the inclusion cutoff.

**Figure S9.** Land-use contrasts across all separate trophic-mode categories.

**Figure S10.** Class-level land-use contrasts for dominant modeled classes.

**Figure S11.** Read coverage of retained FUNGuild assignments.

## Notes

### Competing Interest Statement

The authors have declared no competing interest.

## References

Abarenkov K, Zirk A, Piirmann T, Pöhönen R, Ivanov F, Nilsson RH, Kõljalg U (2023) UNITE general FASTA release for eukaryotes. Version 18.07.2023. UNITE Community. 10.15156/BIO/2938069

Abarenkov K, Nilsson RH, Larsson KH, Taylor AFS, May TW, Frøslev TG et al (2024) The UNITE database for molecular identification and taxonomic communication of fungi and other eukaryotes: sequences, taxa and classifications reconsidered. Nucleic Acids Research 52:D791– D797. 10.1093/nar/gkad1039

Albornoz FE, Ryan MH, Bending GD, Hilton S, Dickie IA, Gleeson DB, Standish RJ (2022) Agricultural land-use favours Mucoromycotinian, but not Glomeromycotinian, arbuscular mycorrhizal fungi across ten biomes. New Phytologist 233:1369–1382. 10.1111/nph.17780

Alguacil MM, Torrecillas E, Lozano Z, Torres MP, Roldán A (2014) Prunus persica crop management differentially promotes arbuscular mycorrhizal fungi diversity in a tropical agro-ecosystem. PLOS ONE 9:e88454. 10.1371/journal.pone.0088454

Anderson MJ (2001) A new method for non-parametric multivariate analysis of variance. Austral Ecology 26:32–46. 10.1111/j.1442-9993.2001.01070.pp.x

Anderson MJ (2006) Distance-Based Tests for Homogeneity of Multivariate Dispersions. Biometrics 62:245–253. 10.1111/j.1541-0420.2005.00440.x

Bahram M, Kõljalg U, Courty PE, Diédhiou AG, Kjøller R, Põlme S, Ryberg M, Veldre V, Tedersoo L (2013) The distance decay of similarity in communities of ectomycorrhizal fungi in different ecosystems and scales. Journal of Ecology 101:1335–1344. 10.1111/1365-2745.12120

Banerjee S, Zhao C, Garland G, Edlinger A, García-Palacios P, Romdhane S, et al (2024) Biotic homogenization, lower soil fungal diversity and fewer rare taxa in arable soils across Europe. Nature Communications 15:327. 10.1038/s41467-023-44073-6

Bardgett RD, van der Putten WH (2014) Belowground biodiversity and ecosystem functioning. Nature 515:505–511. 10.1038/nature13855

Benjamini Y, Hochberg Y (1995) Controlling the False Discovery Rate: A Practical and Powerful Approach to Multiple Testing. Journal of the Royal Statistical Society Series B: Statistical Methodology 57:289–300. 10.1111/j.2517-6161.1995.tb02031.x

Berkelmann D, Schneider D, Meryandini A, Daniel R (2020) Unravelling the effects of tropical land use conversion on the soil microbiome. Environmental Microbiome 15:5. 10.1186/s40793-020-0353-3

Blackwell M (2011) The *Fungi*: 1, 2, 3 … 5.1 million species? American Journal of Botany 98:426–438. 10.3732/ajb.1000298

Brinkmann N, Schneider D, Sahner J, Ballauff J, Edy N, Barus H, et al (2019) Intensive tropical land use massively shifts soil fungal communities. Scientific Reports 9:3403. 10.1038/s41598-019-39829-4

Callahan BJ, McMurdie PJ, Rosen MJ, Han AW, Johnson AJA, Holmes SP (2016) DADA2: High-resolution sample inference from Illumina amplicon data. Nature Methods 13:581–583. 10.1038/nmeth.3869

Chao A, Gotelli NJ, Hsieh TC, Sander EL, Ma KH, Colwell RK, Ellison AM (2014) Rarefaction and extrapolation with Hill numbers: a framework for sampling and estimation in species diversity studies. Ecological Monographs 84:45–67. 10.1890/13-0133.1

Díaz-Vallejo EJ, Seeley M, Smith AP, Marín-Spiotta E (2021) A meta-analysis of tropical land-use change effects on the soil microbiome: Emerging patterns and knowledge gaps. Biotropica 53:738–752. 10.1111/btp.12931

Dumbrell AJ, Ashton PD, Aziz N, Feng G, Nelson M, Dytham C, Fitter AH, Helgason T (2011) Distinct seasonal assemblages of arbuscular mycorrhizal fungi revealed by massively parallel pyrosequencing. New Phytologist 190:794–804. 10.1111/j.1469-8137.2010.03636.x

Goldmann K, Schröter K, Pena R, Schöning I, Schrumpf M, Buscot F, Polle A, Wubet T (2016) Divergent habitat filtering of root and soil fungal communities in temperate beech forests. Scientific Reports 6:31439. 10.1038/srep31439

Gschwend F, Hartmann M, Mayerhofer J, Hug AS, Enkerli J, Gubler A, Meuli RG, Frey B, Widmer F (2022) Site and land-use associations of soil bacteria and fungi define core and indicative taxa. FEMS Microbiology Ecology 97:fiab165. 10.1093/femsec/fiab165

Gumiere T, Durrer A, Bohannan BJM, Andreote FD (2016) Biogeographical patterns in fungal communities from soils cultivated with sugarcane. Journal of Biogeography 43:2016–2026. 10.1111/jbi.12775

Guzman A, Montes M, Hutchins L, DeLaCerda G, Yang P, Kakouridis A, Dahlquist-Willard RM, Firestone MK, Bowles T, Kremen C (2021) Crop diversity enriches arbuscular mycorrhizal fungal communities in an intensive agricultural landscape. New Phytologist 231:447–459. 10.1111/nph.17306

Häkkinen L, Pessi IS, Salonen A-R, Uhlgren O, Soinne H, Hultman J, Heinonsalo J (2025) Fungal communities in boreal soils are influenced by land use, agricultural soil management, and depth. FEMS Microbiology Ecology 101:fiaf002. doi:10.1093/femsec/fiaf002.

Hartmann M, Frey B, Mayer J, Mäder P, Widmer F (2015) Distinct soil microbial diversity under long-term organic and conventional farming. The ISME Journal 9:1177–1194. 10.1038/ismej.2014.210

Hawksworth DL, Lücking R (2017) Fungal Diversity Revisited: 2.2 to 3.8 Million Species. Microbiology Spectrum 5. 10.1128/microbiolspec.funk-0052-2016

Horn S, Hempel S, Verbruggen E, Rillig MC, Caruso T (2017) Linking the community structure of arbuscular mycorrhizal fungi and plants: a story of interdependence? The ISME Journal 11:1400–1411. 10.1038/ismej.2017.5

Hsieh TC, Ma KH, Chao A (2016) iNEXT: an R package for rarefaction and extrapolation of species diversity (Hill numbers). Methods in Ecology and Evolution 7:1451–1456. 10.1111/2041-210x.12613

Keddy PA (1992) Assembly and response rules: two goals for predictive community ecology. Journal of Vegetation Science 3:157–164. 10.2307/3235676

Lee J, Lee S, Young JPW (2008) Improved PCR primers for the detection and identification of arbuscular mycorrhizal fungi. FEMS Microbiology Ecology 65:339–349. 10.1111/j.1574-6941.2008.00531.x

Liu S, Vasar M, Öpik M, Koorem K. (2023) Disturbance induces similar shifts in arbuscular mycorrhizal fungal communities from grassland and arable field soils. Mycorrhiza 33:153–164.

Lumibao CY, Kimbrough ER, Day RH, Conner WH, Krauss KW, Van Bael SA (2020) Divergent biotic and abiotic filtering of root endosphere and rhizosphere soil fungal communities along ecological gradients. FEMS Microbiology Ecology 96:fiaa124. 10.1093/femsec/fiaa124

Manoharan L, Rosenstock NP, Williams A, Hedlund K (2017). Agricultural management practices influence AMF diversity and community composition with cascading effects on plant productivity. Applied Soil Ecology 115:53–59. 10.1016/j.apsoil.2017.03.012

McMurdie PJ, Holmes S (2013) phyloseq: An R package for reproducible interactive analysis and graphics of microbiome census data. PLOS ONE 8:e61217. 10.1371/journal.pone.0061217

Mikryukov V, Dulya O, Zizka A, Bahram M, Hagh-Doust N, Anslan S, et al (2023) Connecting the multiple dimensions of global soil fungal diversity. Science Advances 9:eadj8016. 10.1126/sciadv.adj8016

Nemergut DR, Schmidt SK, Fukami T, O’Neill SP, Bilinski TM, Stanish LF, et al (2013) Patterns and Processes of Microbial Community Assembly. Microbiology and Molecular Biology Reviews 77:342–356. 10.1128/mmbr.00051-12

Nguyen NH, Song Z, Bates ST, Branco S, Tedersoo L, Menke J, Schilling JS, Kennedy PG (2016) FUNGuild: An open annotation tool for parsing fungal community datasets by ecological guild. Fungal Ecology 20:241–248. 10.1016/j.funeco.2015.06.006

Oksanen J, Simpson GL, Blanchet FG, Kindt R, Legendre P, Minchin PR et al (2026) vegan: Community Ecology Package. R package version 2.7–5. 10.32614/CRAN.package.vegan

Özkurt E, Fritscher J, Soranzo N, Ng DYK, Davey RP, Bahram M, Hildebrand F (2022) LotuS2: an ultrafast and highly accurate tool for amplicon sequencing analysis. Microbiome 10:176. 10.1186/s40168-022-01365-1

Palarea-Albaladejo J, Martín-Fernández JA (2015) ZCompositions — R package for multivariate imputation of left-censored data under a compositional approach. Chemometrics and Intelligent Laboratory Systems 143:85–96. 10.1016/j.chemolab.2015.02.019

Powell JR, Rillig MC (2018) Biodiversity of arbuscular mycorrhizal fungi and ecosystem function. New Phytologist 220:1059–1075. 10.1111/nph.15119

Quast C, Pruesse E, Yilmaz P, Gerken J, Schweer T, Yarza P, Peplies J, Glöckner FO (2013) The SILVA ribosomal RNA gene database project: improved data processing and web-based tools. Nucleic Acids Research 41:D590–D596. 10.1093/nar/gks1219

Rognes T, Flouri T, Nichols B, Quince C, Mahé F (2016) VSEARCH: a versatile open source tool for metagenomics. PeerJ 4:e2584. 10.7717/peerj.2584

Rousset F, Ferdy JB (2014) Testing environmental and genetic effects in the presence of spatial autocorrelation. Ecography 37:781–790. 10.1111/ecog.00566

Saravia-Maldonado SA, Ramírez-Rosario B, Rodríguez-González MÁ, Fernández-Pozo LF (2025) Land Use Changes Influence Tropical Soil Diversity: An Assessment Using Soil Taxonomy and the World Reference Base for Soil Classifications. Agriculture 15:1893. 10.3390/agriculture15171893

Song H, Singh D, Tomlinson KW, Yang X, Ogwu MC, Slik JWF, Adams JM (2019) Tropical forest conversion to rubber plantation in southwest China results in lower fungal beta diversity and reduced network complexity. FEMS Microbiology Ecology 95:fiz092. 10.1093/femsec/fiz092

Stewart JD, Ramirez DX, Leon-Reyes A, Barriga N, Llerena S, Manley BF, Carpintero-Salvador N, Ruiz-Uriguen M, Raaijmakers JM, Kiers ET, Weedon JT (2026) Land Use Change Reshapes Climate-Driven Diversity Patterns of Tropical Arbuscular Mycorrhizal Fungi. Molecular Ecology 35:e70253. doi:10.1111/mec.70253.

Tedersoo L, Bahram M, Põlme S, Kõljalg U, Yorou NS, Wijesundera R, et al (2014) Global diversity and geography of soil fungi. Science 346:1256688. 10.1126/science.1256688

Tedersoo L, Anslan S, Bahram M, Drenkhan R, Pritsch K, Buegger F, et al (2020) Regional-Scale In-Depth Analysis of Soil Fungal Diversity Reveals Strong pH and Plant Species Effects in Northern Europe. Frontiers in Microbiology 11:1953. 10.3389/fmicb.2020.01953

Tedersoo L, Bahram M, Zinger L, Nilsson RH, Kennedy PG, Yang T, Anslan S, Mikryukov V (2022) Best practices in metabarcoding of fungi: From experimental design to results. Molecular Ecology 31:2769–2795. 10.1111/mec.16460

Tsiafouli MA, Thébault E, Sgardelis SP, de Ruiter PC, van der Putten WH, Birkhofer K, et al (2015) Intensive agriculture reduces soil biodiversity across Europe. Global Change Biology 21:973–985. 10.1111/gcb.12752

Tuck SL, Winqvist C, Mota F, Ahnström J, Turnbull LA, Bengtsson J (2014) Land-use intensity and the effects of organic farming on biodiversity: a hierarchical meta-analysis. Journal of Applied Ecology 51:746–755. 10.1111/1365-2664.12219

Vályi K, Mardhiah U, Rillig MC, Hempel S (2016) Community assembly and coexistence in communities of arbuscular mycorrhizal fungi. The ISME Journal 10:2341–2351. 10.1038/ismej.2016.46

van der Heijden MGA, Klironomos JN, Ursic M, Moutoglis P, Streitwolf-Engel R, Boller T, Wiemken A, Sanders IR (1998) Mycorrhizal fungal diversity determines plant biodiversity, ecosystem variability and productivity. Nature 396:69–72. 10.1038/23932

van Rijssel SQ, Veen GFC, Koorneef GJ, Bakx-Schotman JMTT, Ten Hooven FC, Geisen S, van der Putten WH (2022) Soil microbial diversity and community composition during conversion from conventional to organic agriculture. Molecular Ecology 31:4017–4030. doi:10.1111/mec.16571.

Vellend M (2010) Conceptual Synthesis in Community Ecology. The Quarterly Review of Biology 85:183–206. 10.1086/652373

Wagg C, Bender SF, Widmer F, van der Heijden MGA (2014) Soil biodiversity and soil community composition determine ecosystem multifunctionality. Proceedings of the National Academy of Sciences 111:5266–5270. 10.1073/pnas.1320054111

Wang M, Zhao Y, Zhang B (2015) Efficient Test and Visualization of Multi-Set Intersections. Scientific Reports 5:16923. 10.1038/srep16923

White TJ, Bruns T, Lee S, Taylor J (1990) Amplification and direct sequencing of fungal ribosomal RNA genes for phylogenetics. In: PCR protocols: a guide to methods and applications, pp315–322. 10.1016/B978-0-12-372180-8.50042-1

