## Supplementary material for "Agricultural Management Restructures the Functional Composition of Soil Fungal Communities Across Tropical Landscapes": Electronic Supplemental Resource 2

**Online Resource 2**

### Supplemental Note S1: Bioinformatic processing of ITS2 and SSU libraries

Land use and geography structure soil fungal communities in Honduras

Evelin Y. Reyes-Mendez; Cleber Ten Caten; Pedro Henrique Dias dos Santos; David Galo; Daijiang Li; Jordan A. Dowell

Corresponding author: Jordan A. Dowell

Department of Biological Sciences, Louisiana State University, Baton Rouge, Louisiana 70802, USA

#### Scope and source records

This note documents the original processing of the ITS2 and small-subunit ribosomal RNA (SSU) libraries using archived LotuS2 run logs, generated command logs, input mapping files, and SDM quality-filter configurations. LotuS2 reported version 2.34; its recorded package-directory identifier was 2.34.1-0 (Özkurt et al. 2022).

The original pipeline used paired-end files from two sequencing runs. Mapping files contain 124 sample identifiers for each marker, FASTQ basenames, sequencing-run labels, and primer sequences. ITS2 primers were GCATCGATGAAGAACGCAGC and TCCTCCGCTTATTGATATGC; SSU primers were CAGCCGCGGTAATTCCAGCT and GAACCCAAACACTTTGGTTTCC. Primer sequences were supplied through the mapping files rather than explicit primer arguments in the logged LotuS2 commands. Archived processing metadata are retained for provenance; the reconciled analytical metadata are in Table S1.

The logs label the processing profile as MiSeq and use MiSeq-named SDM configuration files. These identify the filtering profile; the physical sequencing instrument is reported separately in the manuscript.

#### Path-normalized pipeline commands

The commands below retain every non-path argument in the logged top-level calls. Original host-specific paths have been replaced by the aliases listed here, and copied input mapping files have descriptive filenames. Set the aliases to the local executable, library, mapping, configuration, and reference locations before running these commands. The two comma-separated SSU reference lists preserve their recorded order. The commands are a record of the original processing settings; recovering the original reference FASTA and taxonomy files is required for a complete sequence-processing rerun.

**LOTUS2_EXEC:** Installed LotuS2 version 2.34 Perl executable

**LIBRARY_DIR:** Directory holding the paired-end library FASTQ files

**OUTPUT_DIR:** New output directory for the processing run

**MAPPING_DIR:** Directory holding ITS2_processing_in.map and SSU_processing_in.map

**CONFIG_DIR:** Directory holding the two archived SDM configuration files

**VT_FASTA:** vt_types_fasta_from_05-06-2019.qiime.fasta

**VT_TAX:** vt_types_GF.txt

**SILVA_FASTA:** SLV_138.1_SSU_NO_AMF.fasta

**SILVA_TAX:** SLV_138.1_SSU_NO_AMF.tax

##### ITS2 OTU processing

```
perl "$LOTUS2_EXEC" \
  -i "$LIBRARY_DIR" -o "$OUTPUT_DIR/ITS2" \
  -m "$MAPPING_DIR/ITS2_processing_in.map" \
  -refDB UNITE -amplicon_type ITS2 \
  -LCA_idthresh 97,95,93,91,88,78 \
  -tax_group fungi -taxAligner blast \
  -clustering vsearch -derepMin 10:1,5:2,3:3 \
  -sdmopt "$CONFIG_DIR/sdm_miSeq_ITS.txt" -id 0.97
```

##### SSU ASV processing

```
perl "$LOTUS2_EXEC" \
  -i "$LIBRARY_DIR" -o "$OUTPUT_DIR/SSU" \
  -m "$MAPPING_DIR/SSU_processing_in.map" \
  -refDB "$VT_FASTA,$SILVA_FASTA" \
  -tax4refDB "$VT_TAX,$SILVA_TAX" \
  -amplicon_type SSU -LCA_idthresh 97,95,93,91,88,78,0 \
  -tax_group fungi -taxAligner blast -clustering dada2 \
  -LCA_cover 0.97 -derepMin 10:1,5:2,3:3 \
  -sdmopt "$CONFIG_DIR/sdm_miSeq2.txt"
```

#### Read filtering and sequence inference

SDM performed demultiplexing, primer/barcode removal, quality filtering, and dereplication. All 30 non-comment options in each archived configuration are reproduced in the parameter table, including their original spelling. Sequence-length settings refer to processed reads after primer/barcode removal and trimming. Asterisk-prefixed settings define an alternative read stream used for abundance recovery rather than the initial construction of sequence representatives. Both primary streams required mean quality 27 and zero ambiguous nucleotides. Alternative streams required mean quality 20, allowed one ambiguous nucleotide, and disabled the accumulated-error trimming threshold. Their minimum lengths were 110 nt for ITS2 and 170 nt for SSU.

ITS2 primary reads had a minimum length of 110 nt, a maximum length of 1,000 nt, no fixed-length truncation, and an accumulated-error threshold of 0.75. Both forward and reverse primer matches were required, with zero primer mismatches. SSU primary reads had a minimum length of 100 nt, fixed truncation at 170 nt, removal of the initial 5 nt, and an accumulated-error threshold of 3.75. A forward primer match was required; a reverse primer match was optional. Shared quality-window, end-trimming, homopolymer, and probabilistic-error settings are listed in the table. These processing lengths do not establish the lengths of the final extended representative sequences.

Both markers used the recorded dereplication settings 10:1,5:2,3:3. ITS2 operational taxonomic units (OTUs) were clustered with VSEARCH 2.15.0 at 97% identity (Rognes et al. 2016). SSU amplicon sequence variants (ASVs) were inferred with DADA2 through the LoTS2 pipeline (Callahan et al. 2016), with the logged error-profile setting -dada2seed 0. The pipeline did not merge read pairs before initial clustering/inference. It subsequently extended sequence representatives and merged their paired reads. Additional dereplicated and alternative-quality reads were recovered against representatives using minimap2 at 97% identity for ITS2 and 99% for SSU (Li 2018). The recorded abundance-mapping coverage criterion was 0.8.

**All non-comment SDM options are preserved from the archived configuration files**

| Recorded option | ITS2 | SSU | Interpretation |
|---|---|---|---|
| minSeqLength | 110 | 100 | Minimum processed sequence length(nt) |
| maxSeqLength | 1000 | 1000 | Maximum processed sequence length(nt) |
| minAvgQuality | 27 | 27 | Minimum mean quality score |
| *minSeqLength | 110 | 170 | Alternative-stream minimum length(nt) |
| *minAvgQuality | 20 | 20 | Alternative-stream minimum mean quality |
| TruncateSequenceLength | -1 | 170 | Fixed processed length;−1 disables truncation |
| maxAmbiguousNT | 0 | 0 | Maximum ambiguous nucleotides |

| Recorded option | ITS2 | SSU | Interpretation |
|---|---|---|---|
| *maxAmbiguousNT | 1 | 1 | Alternative-stream maximum ambiguous nucleotides |
| maxHomonucleotide | 8 | 8 | Maximum homopolymer run length |
| QualWindowWidth | 50 | 50 | Quality-filter window width(nt) |
| QualWindowThreshhold | 25 | 25 | Mean-quality threshold for that window |
| TrimWindowWidth | 20 | 20 | End-trimming window width(nt) |
| TrimWindowThreshhold | 25 | 25 | Mean-quality threshold for end trimming |
| maxAccumulatedError | 0.75 | 3.75 | Accumulated-error threshold for trimming |
| *maxAccumulatedError | -1 | -1 | Alternative-stream threshold;−1 disables it |
| BinErrorModelMaxExpError | 2.5 | 2.5 | Binomial expected-error parameter |
| BinErrorModelAlpha | 0.005 | 0.005 | Binomial model alpha parameter |
| maxBarcodeErrs | 0 | 0 | Maximum barcode mismatches |
| maxPrimerErrs | 0 | 0 | Maximum primer mismatches |
| keepBarcodeSeq | 0 | 0 | Retain barcode sequence(0=no) |
| keepPrimerSeq | 0 | 0 | Retain primer sequence(0=no) |
| fastqVersion | auto | auto | Quality-score encoding detection |
| TechnicalAdapter | (blank) | (blank) | Additional technical adapter sequence |
| TrimStartNTs | 0 | 5 | Number of initial nucleotides removed |
| PEheaderPairFmt | 1 | 1 | Paired-end header-format option |
| RejectSeqWithoutRevPrim | T | F | Reject reads without reverse-primer match |
| RejectSeqWithoutFwdPrim | T | T | Reject reads without forward-primer match |
| AmpliconShortPE | T | T | Short paired-end amplicon handling |
| CheckForMixedPairs | T | T | Check for switched pairs |
| CheckForReversedSeqs | T | T | Check reversed sequence orientation |

T=true;F=false;0=no for retention options. −1 disables the documented setting. (blank) records an empty parameter value. Asterisk-prefixed options apply to the alternative abundance-recovery stream. Original option spelling, including Threshhold and Homonucleotide, is retained.

#### Chimera and target-marker processing

ITS2 command logs show VSEARCH uchime_denovo with abundance-skew parameter 2 and uchime_ref with the packaged reference uchime_sh_refs_dynamic_develop_985_11.03.2015.ITS2.fasta (Nilsson et al. 2015; Rognes et al. 2016). The run log reports zero representatives removed at the de novo stage and 311 at the reference-filter stage; its final matrix summary reports 272 chimeric feature entries representing 202,711 reads. These counts describe different processing stages. ITSx subsequently detected ITS regions and removed representatives failing its target-region processing (Bengtsson-Palme et al. 2013). The command requested ITS1 and ITS2 regions, partial 0 and preservation of sequence identifiers; the executable-directory identifier was ITSx_1.1.4. PhiX screening with minimap2 reported zero contaminated features.

For SSU, the executed pipeline states No ref based chimera detection; its generated command log contains no reference-based or separate VSEARCH de novo chimera-filter command. Consequently, the generic configuration-header entries for chimera checking do not establish that those separate filters were executed. DADA2 inference and its reported outputs are documented by the archived pipeline logs. PhiX screening reported zero contaminated features.

#### Taxonomic assignment and reference provenance

ITS2 representatives were searched against UNITE v9 reference file sh_refs_v9_25.07.2023.fasta with its paired taxonomy file sh_refs_v9_25.07.2023.tax. The archived pipeline citation file identifies the UNITE general FASTA release for eukaryotes, DOI 10.15156/BIO/2938069 (Abarenkov et al. 2023); the UNITE repository identifies that dataset as version 18.07.2023. Abarenkov et al. (2024) describe the UNITE database methods.

SSU classification used two custom references in the recorded order: vt_types_fasta_from_05-06-2019.qiime.fasta with vt_types_GF.txt, followed by SLV_138.1_SSU_NO_AMF.fasta with SLV_138.1_SSU_NO_AMF.tax.

Both markers were analyzed using BLASTn with up to 200 target sequences per query, a 75% search identity floor, searches on both strands, and eight threads (Altschul et al. 1990). The LotuS2 LCA step used fraction 0.8, minimum alignment length 70, and identity-threshold vector 97,95,93,91,88,78,0. Alignment coverage was 0.5 for ITS2 and 0.97 for SSU. The ITS2 top-level command lists 97,95,93,91,88,78; its executed LCA command appends the final 0. These LCA settings are taxonomic-assignment parameters rather than sample read-depth thresholds. *Unclassified* features remained in the pipeline output matrices.

#### Archived outputs and reproducibility scope

The pipeline generated feature-count and taxonomic tables, representative sequences, phyloseq objects, and phylogenetic outputs. Generated commands used MAFFT followed by FastTreeMP with nucleotide/GTR settings (Katoh et al. 2002; Price et al. 2010), then imported the tables into phyloseq (McMurdie and Holmes 2013). The run-log output summaries contain 9,405 ITS2 OTUs and 3,961 SSU ASVs before the manuscript-specific fungal and sample-depth filtering. These pipeline feature counts are distinct from the retained fungal counts used in the revised analyses.

#### References


Abarenkov K, Nilsson RH, Larsson KH, Taylor AFS, May TW, Frøslev TG et al (2024) The UNITE database for molecular identification and taxonomic communication of fungi and other eukaryotes: sequences, taxa and classifications reconsidered. Nucleic Acids Research 52:D791–D797. https://doi.org/10.1093/nar/gkad1039

Abarenkov K, Zirk A, Piirmann T, Pöhönen R, Ivanov F, Nilsson RH, Kõljalg U (2023) UNITE general FASTA release for eukaryotes. Version 18.07.2023. UNITE Community. https://doi.org/10.15156/BIO/2938069

Altschul SF, Gish W, Miller W, Myers EW, Lipman DJ (1990) Basic local alignment search tool. Journal of Molecular Biology 215:403–410. https://doi.org/10.1016/s0022-2836(05)80360-2

Bengtsson-Palme J, Ryberg M, Hartmann M, Branco S, Wang Z, Godhe A, et al (2013) Improved software detection and extraction of ITS1 and ITS2 from ribosomal ITS sequences of fungi and other eukaryotes for analysis of environmental sequencing data. Methods in Ecology and Evolution 4:914–919. https://doi.org/10.1111/2041-210x.12073

Callahan BJ, McMurdie PJ, Rosen MJ, Han AW, Johnson AJA, Holmes SP (2016) DADA2: High-resolution sample inference from Illumina amplicon data. Nature Methods 13:581–583. https://doi.org/10.1038/nmeth.3869

Katoh K, Misawa K, Kuma K, Miyata T (2002) MAFFT: a novel method for rapid multiple sequence alignment based on fast Fourier transform. Nucleic Acids Research 30:3059–3066. https://doi.org/10.1093/nar/gkf436

Li H (2018) Minimap2: pairwise alignment for nucleotide sequences. Bioinformatics 34:3094–3100. https://doi.org/10.1093/bioinformatics/bty191

McMurdie PJ, Holmes S (2013) phyloseq: An R package for reproducible interactive analysis and graphics of microbiome census data. PLOS ONE 8:e61217. https://doi.org/10.1371/journal.pone.0061217

Nilsson RH, Tedersoo L, Ryberg M, Kristiansson E, Hartmann M, Unterseher M, et al (2015) A Comprehensive, Automatically Updated Fungal ITS Sequence Dataset for Reference-Based Chimera Control in Environmental Sequencing Efforts. Microbes and environments 30:145–150. https://doi.org/10.1264/jsme2.me14121

Öpik M, Vanatoa A, Vanatoa E, Moora M, Davison J, Kalwij JM, Reier Ü, Zobel M (2010) The online database MaarjAM reveals global and ecosystemic distribution patterns in arbuscular mycorrhizal fungi (*Glomeromycota*). New Phytologist 188:223–241. https://doi.org/10.1111/j.1469-8137.2010.03334.x

Özkurt E, Fritscher J, Soranzo N, Ng DYK, Davey RP, Bahram M, Hildebrand F (2022) LotuS2: an ultrafast and highly accurate tool for amplicon sequencing analysis. Microbiome 10:176. https://doi.org/10.1186/s40168-022-01365-1

Price MN, Dehal PS, Arkin AP (2010) FastTree 2 – Approximately Maximum-Likelihood Trees for Large Alignments. PLoS ONE 5:e9490. https://doi.org/10.1371/journal.pone.0009490

Quast C, Pruesse E, Yilmaz P, Gerken J, Schweer T, Yarza P, Peplies J, Glöckner FO (2013) The SILVA ribosomal RNA gene database project: improved data processing and web-based tools. Nucleic Acids Research 41:D590–D596. https://doi.org/10.1093/nar/gks1219

Rognes T, Flouri T, Nichols B, Quince C, Mahé F (2016) VSEARCH: a versatile open source tool for metagenomics. PeerJ 4:e2584. https://doi.org/10.7717/peerj.2584
