## Supplementary material for "Agricultural Management Restructures the Functional Composition of Soil Fungal Communities Across Tropical Landscapes": Electronic Supplemental Resource 3

Land use and geography structure soil fungal communities in Honduras

Figures S1–S11 and their legends

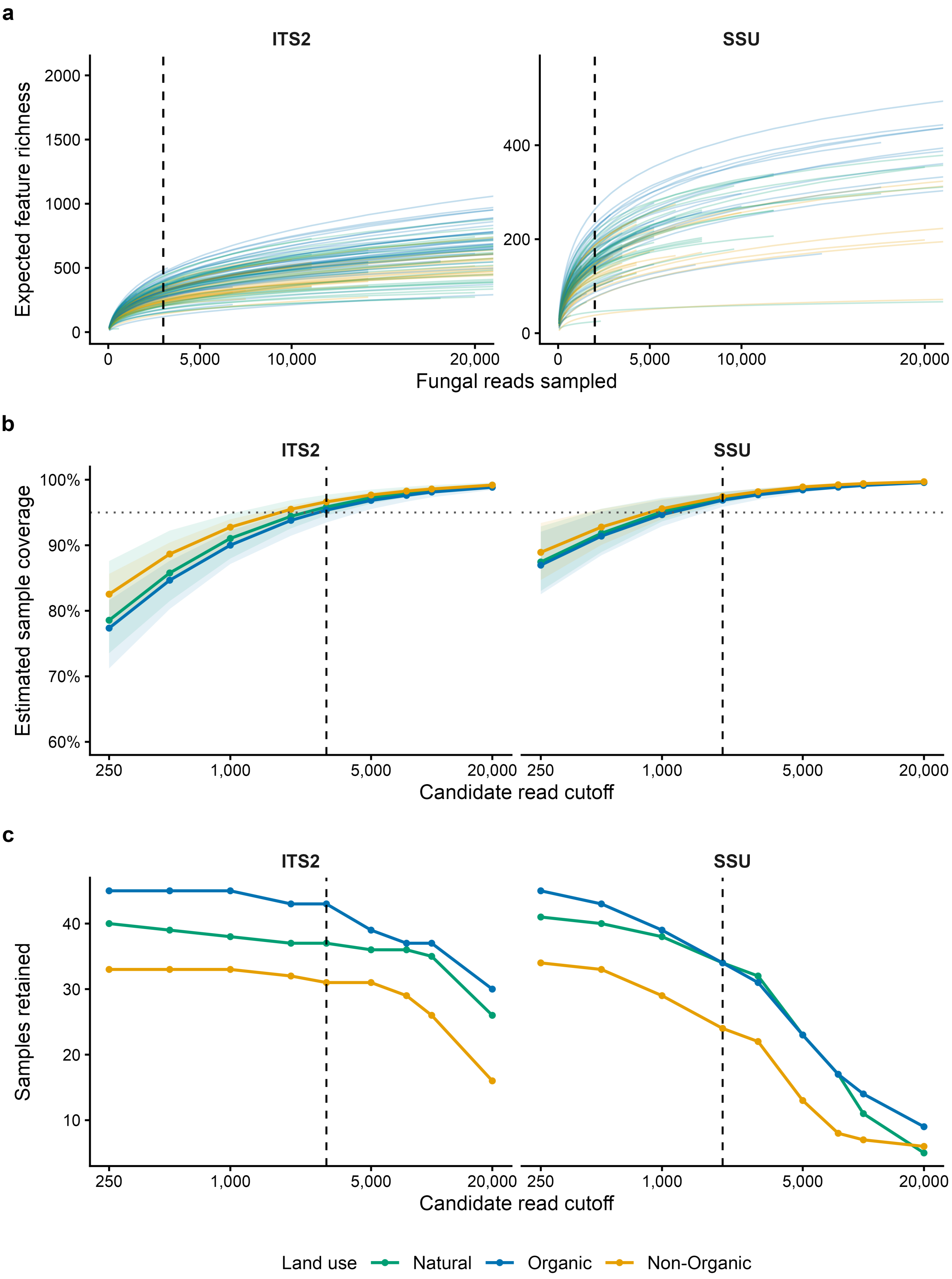


**Fig. S1** Rarefaction curves and read-cutoff selection. (a) Deterministic individual-sample rarefaction curves, shown over the 0-20,000-read range to emphasize the evaluated cutoffs. (b) Median estimated sample coverage by land use at each candidate cutoff, with ribbons showing the 10th-90th percentile range; the dotted line marks 95% coverage. (c) Sample retention by land use across candidate cutoffs. Dashed lines mark the selected marker-specific cutoffs. Sample coverage describes estimated read-probability mass, not the fraction of all taxa recovered. Curves continued to accumulate features at higher depths. Candidate values and the selection rule are reported in Table S1. Selected cutoffs are 3,000 fungal reads for ITS2 and 2,000 for SSU

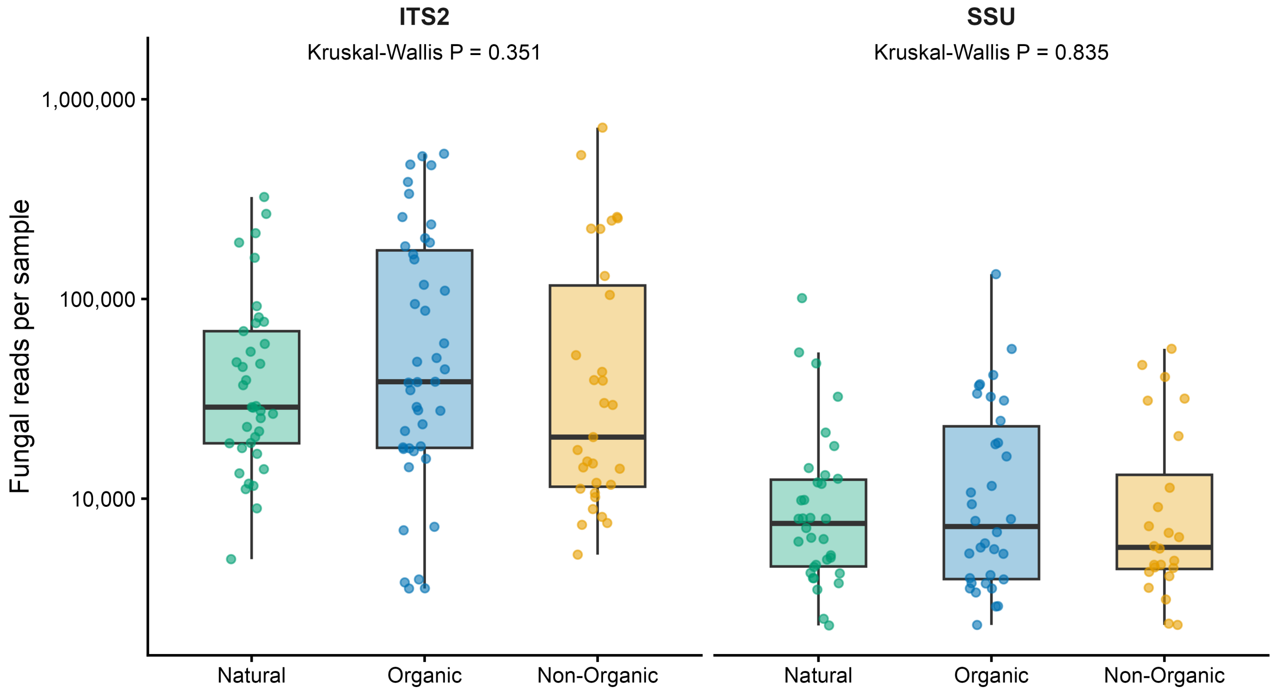


**Fig. S2** Fungal sequencing depths of retained samples across land uses. The primary cutoffs are 3,000 fungal reads for ITS2 and 2,000 for SSU. Points represent individual samples. Boxplots show medians, interquartile ranges, and 1.5-times-interquartile-range whiskers. The read-depth axis is logarithmic. Each panel reports the overall Kruskal-Wallis P value comparing the three land uses. These are overall tests, not pairwise significance labels. Group sample sizes and read-depth tests are reported in Table S1

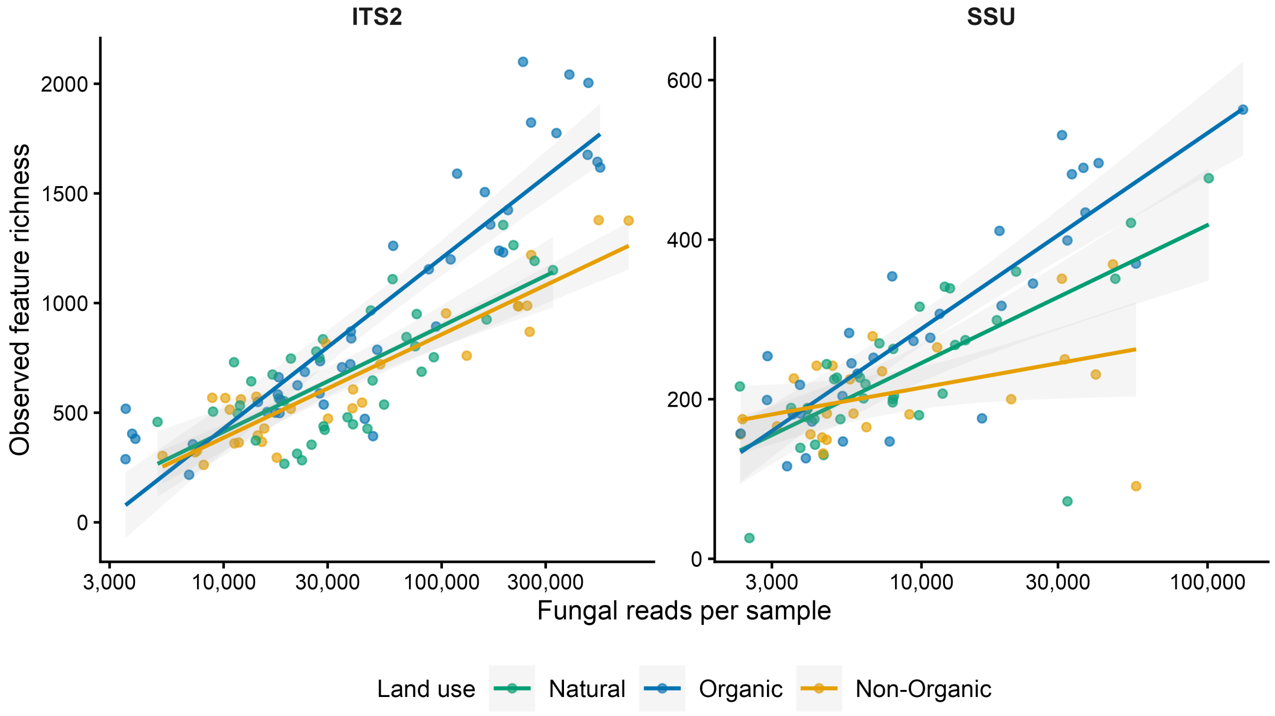


**Fig. S3** Observed richness in relation to fungal sequencing depth. Points represent retained individual samples. Lines and shaded bands show land-use-specific diagnostic linear fits and 95% confidence intervals on log10 read depth. Primary inferential richness models use spatial negative-binomial distributions and include depth-by-land-use interactions. Contrasts are reported in Table S5 and coefficients in Table S8

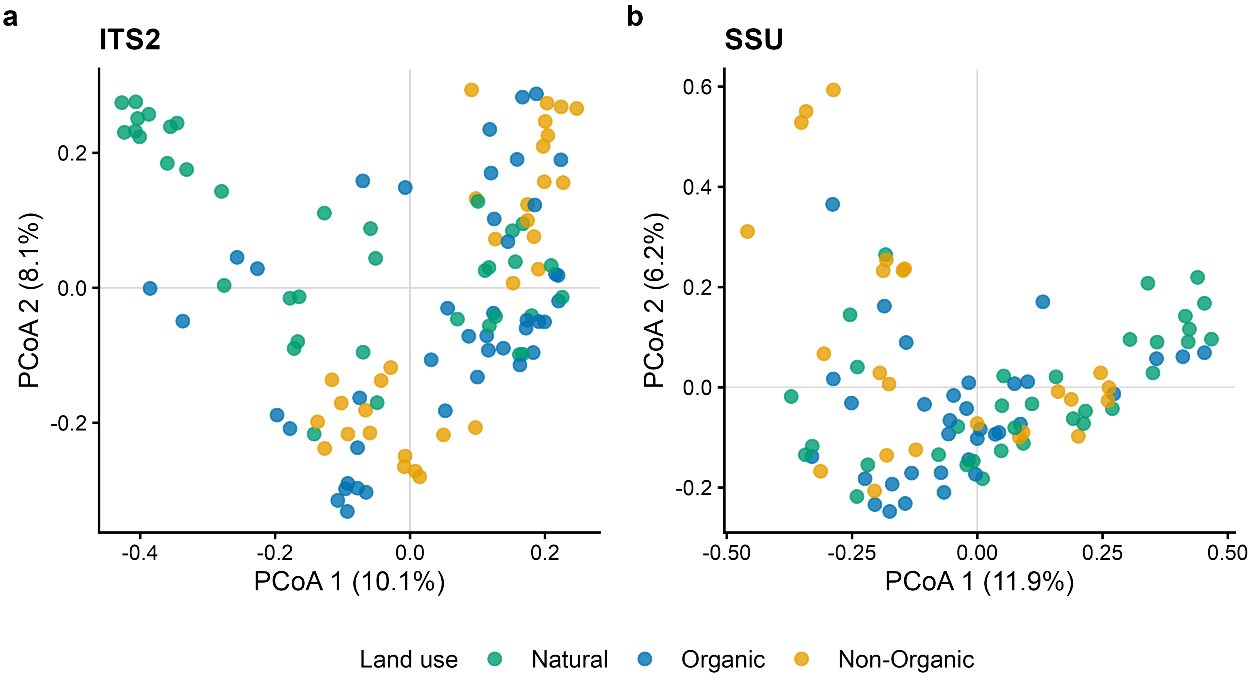


**Fig. S4** Principal coordinates ordination of retained fungal communities. Ordinations use Bray-Curtis dissimilarities calculated from relative fungal read abundances. Points are samples colored by land use. Axis labels give percentages of the positive-eigenvalue sum from the corrected principal coordinates decomposition. Ordinations are descriptive; adjusted community tests are reported in Table S2

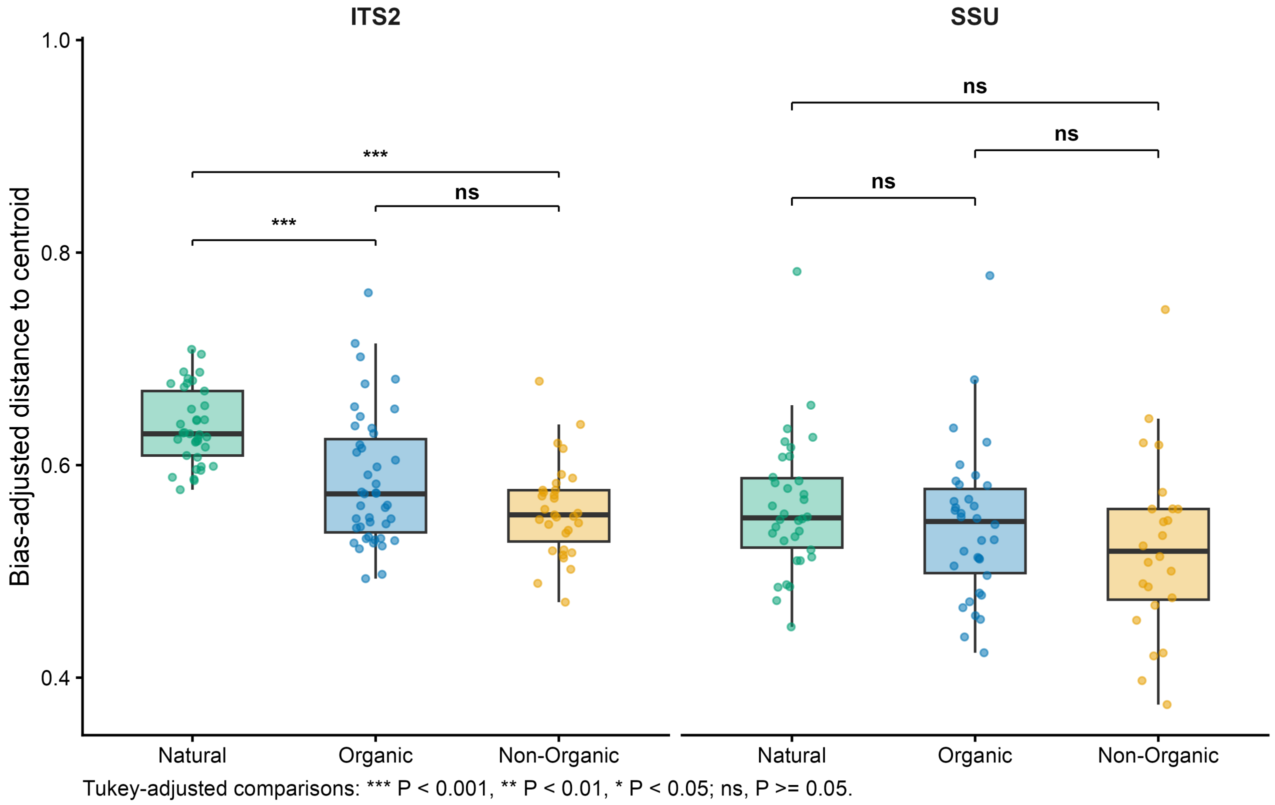


**Fig. S5** Within-land-use community dispersion. Distances to land-use centroids are calculated from Bray-Curtis dissimilarities and adjusted for unequal group sizes. Points are samples, and boxplots show medians, interquartile ranges, and 1.5-times-interquartile-range whiskers. Brackets show Tukey-adjusted pairwise land-use comparisons (* adjusted $P < 0.05$, ** adjusted $P < 0.01$, *** adjusted $P < 0.001$; ns, adjusted $P \geq 0.05$). Overall permutation dispersion tests and exact Tukey-adjusted pairwise comparisons are reported in Table S2

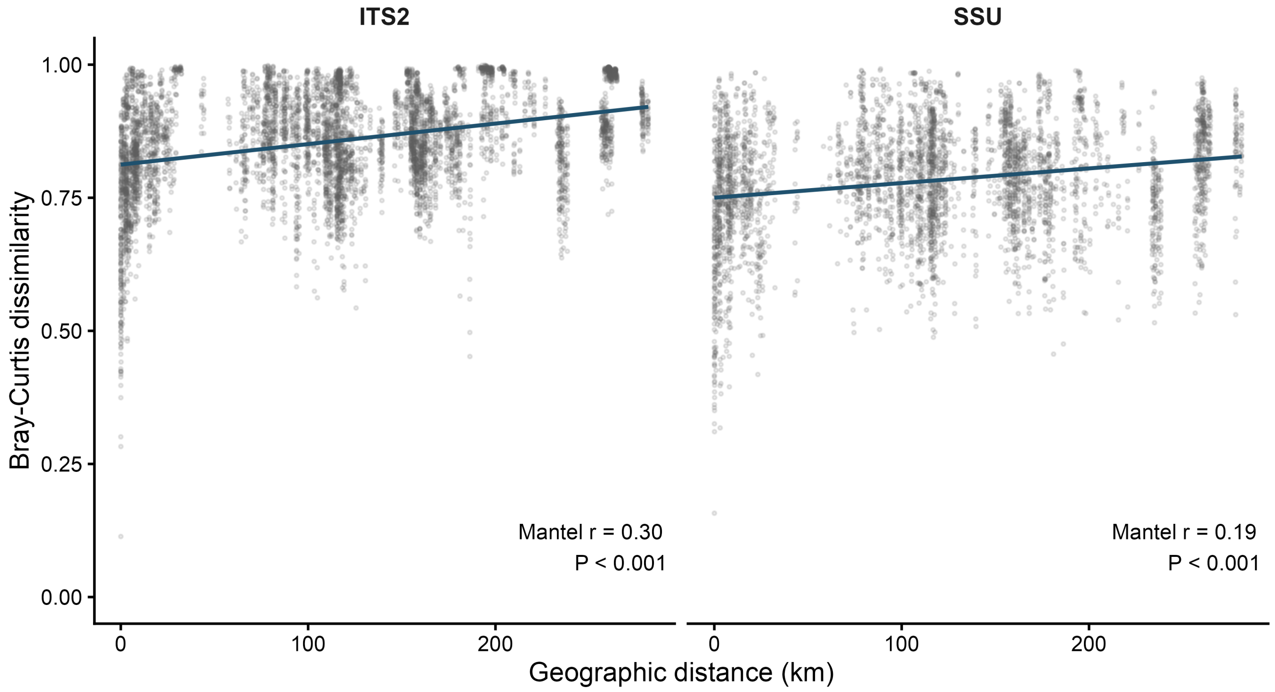


**Fig. S6** Geographic distance-decay across all retained samples. Pairwise Bray-Curtis dissimilarities are plotted against Haversine distances separately for each marker. Lines are descriptive linear trends; labels report Spearman Mantel correlations and 9,999-permutation P values. These correlations are distinct from PERMANOVA proportions of variation explained

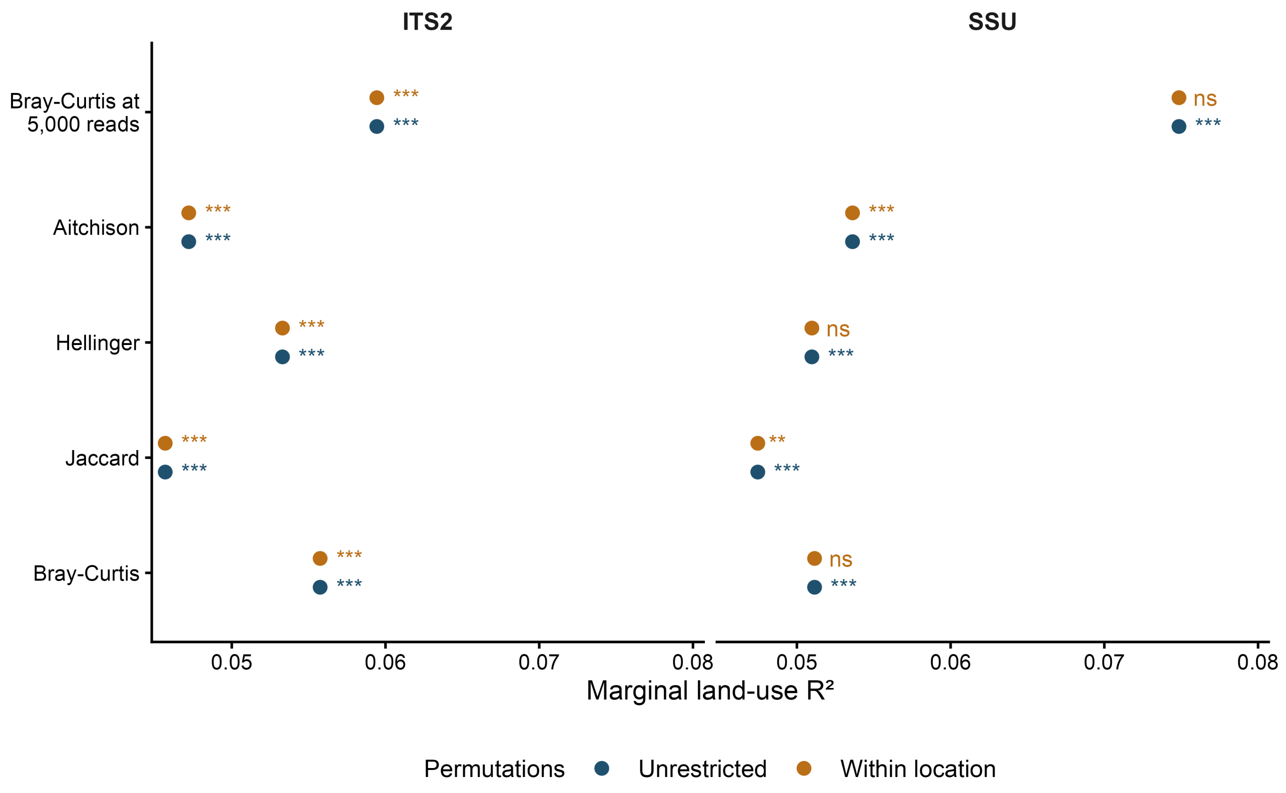


**Fig. S7** Sensitivity of community-composition results to distance metric and read cutoff. Points show marginal PERMANOVA land-use $R^2$ after adjustment for geography, sequencing run, and $\log_{10}$ fungal read depth. The primary dataset is evaluated using Bray-Curtis, Jaccard, Hellinger, and Aitchison distances; Bray-Curtis is additionally evaluated at 5,000 reads. Symbols show unadjusted permutation significance (* $P < 0.05$, ** $P < 0.01$, *** $P < 0.001$; ns, $P \geq 0.05$). Unrestricted and within-location permutations are shown separately. Table S3 reports exact tests, sample counts, and associated dispersion results

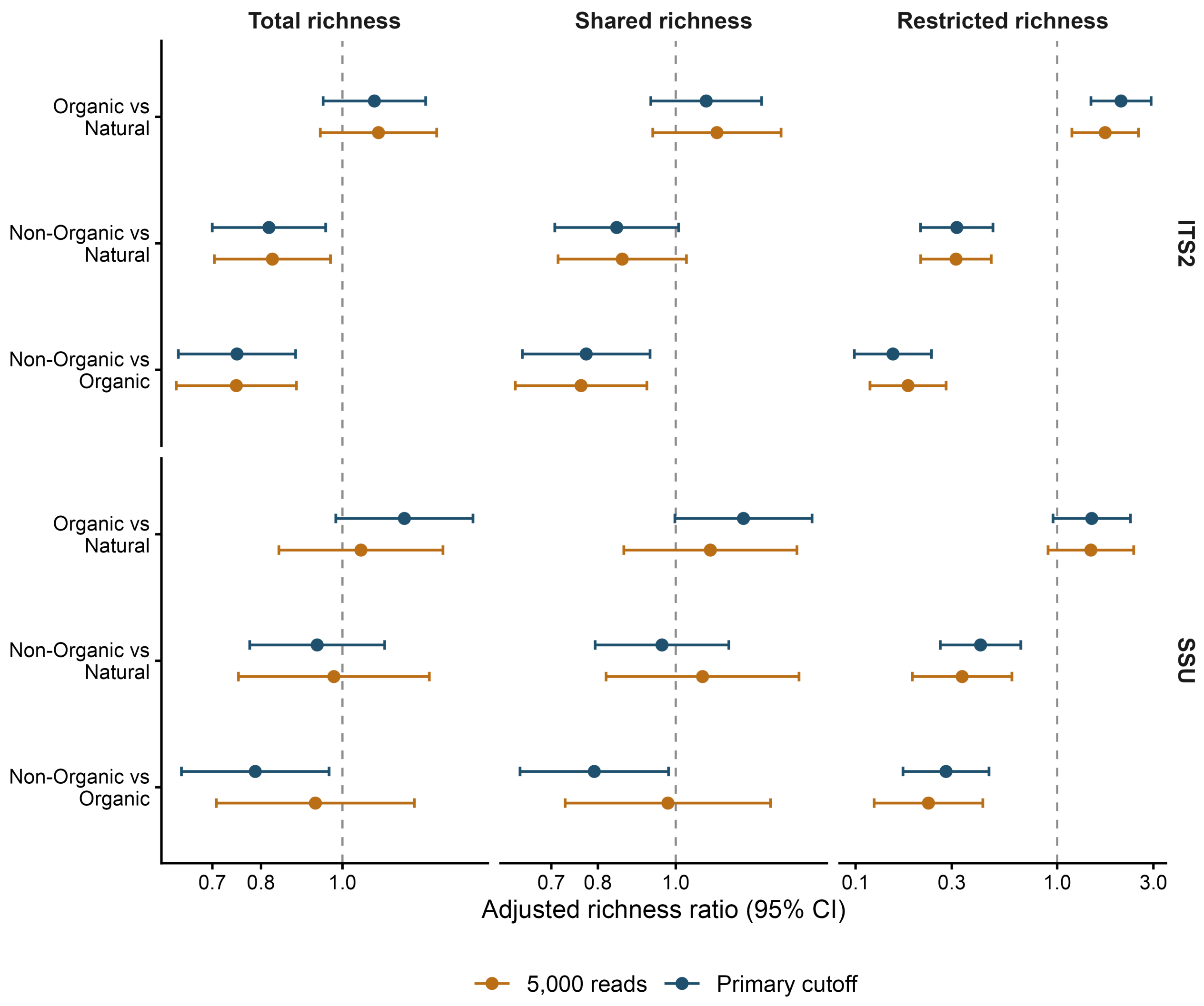


**Fig. S8** Sensitivity of adjusted richness contrasts to the inclusion cutoff. Points and horizontal intervals show model-estimated richness ratios and 95% confidence intervals at the same marker-specific reference read depth for the primary and 5,000-read analyses. The reference depth is the back-transformed median $\log_{10}$ fungal-read depth among samples retained under the primary cutoff. Models include land-use-specific depth slopes, sequencing run, and spatial covariance. The dashed line denotes equal fitted richness. Table S5 reports the primary and 5,000-read richness contrasts

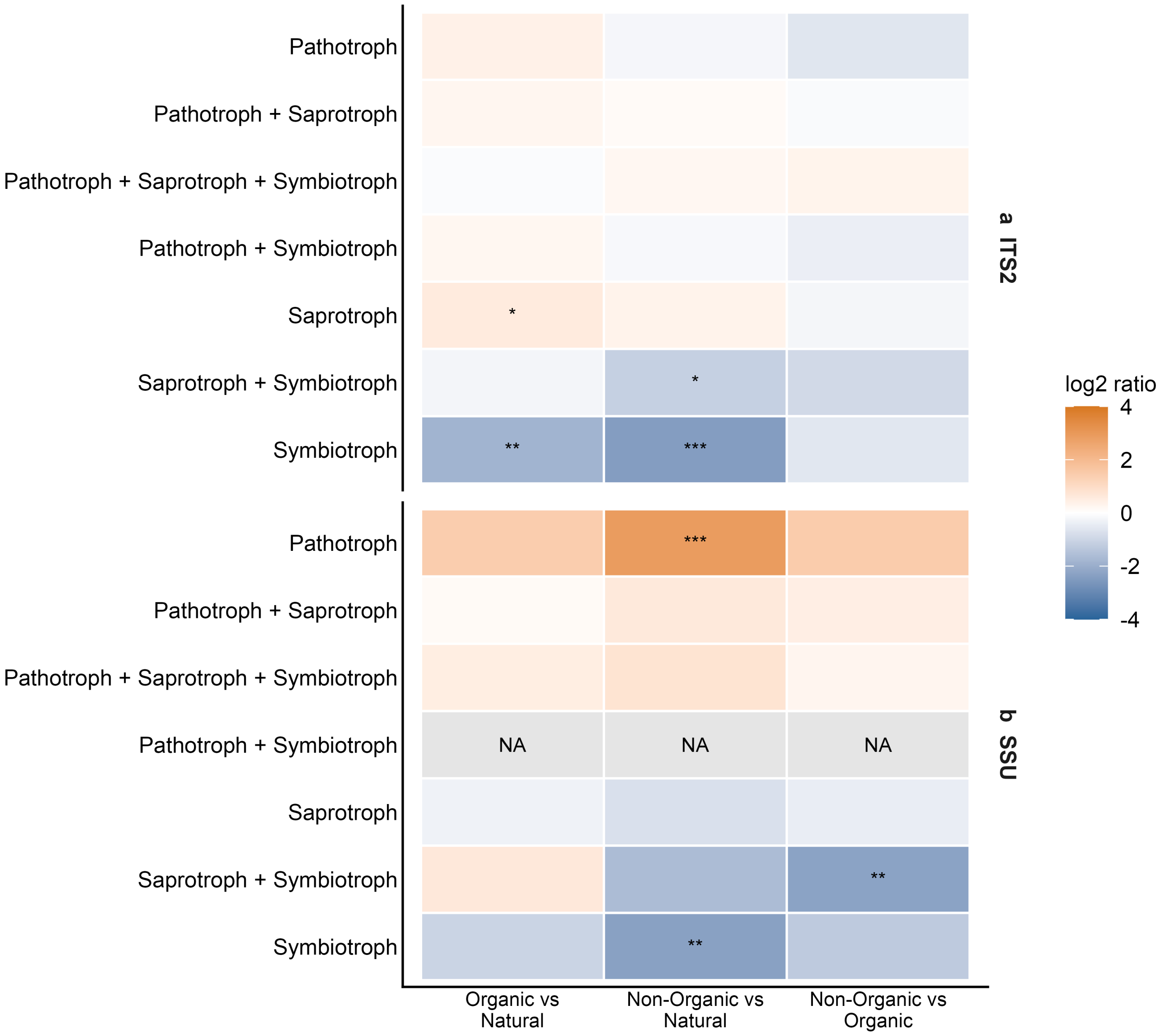


**Fig. S9** Land-use contrasts across all separate trophic-mode categories. Colors show modeled $log_2$ read-abundance ratios, and stars use FDR adjustment across all estimable trophic-mode contrasts within each marker. Grey cells labeled NA indicate a category that did not meet count-model eligibility criteria, not evidence of no effect. Full numerical and screening results are reported in Tables S6 and S7. Marker rows are (a) ITS2 and (b) SSU

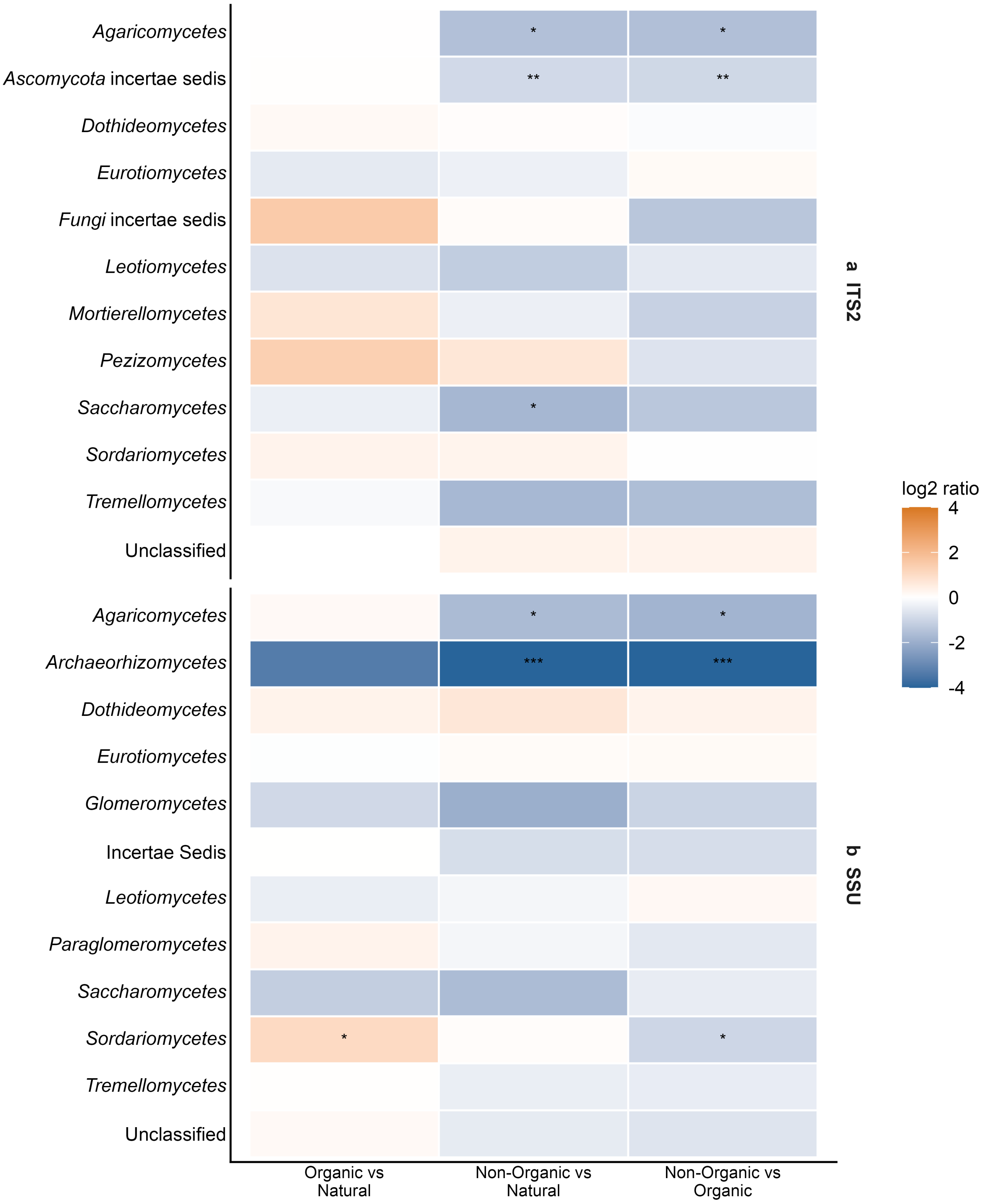


**Fig. S10** Class-level land-use contrasts for dominant modeled classes. Classes are selected by mean relative read abundance within each marker, up to 12 per marker. Color and significance conventions match Figure 4, with FDR adjustment across all eligible class contrasts rather than only the displayed classes. All modeled classes and screening outcomes are available in Tables S6 and S7. Marker rows are (a) ITS2 and (b) SSU

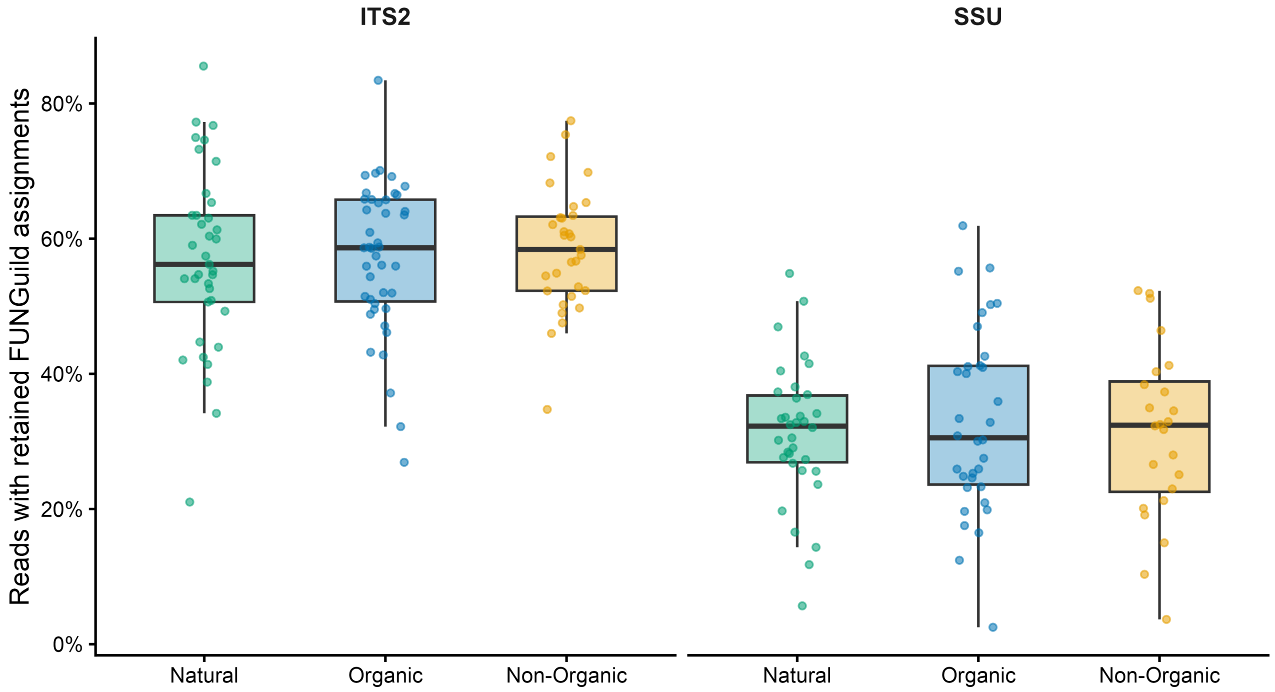


**Fig. S11** Read coverage of retained FUNGuild assignments. The response is the proportion of total fungal reads assigned to a canonical trophic mode with Probable or Highly Probable confidence. Points are retained samples, and boxplots summarize distributions by land use and marker. These boxplots are descriptive; no pairwise hypothesis tests are assigned to these panels. Table S4 reports classification coverage and annotation cleaning
